# Glycolytic flux in *Saccharomyces cerevisiae* is dependent on RNA polymerase III and its negative regulator Maf1

**DOI:** 10.1101/372946

**Authors:** Róża Szatkowska, Manuel Garcia-Albornoz, Katarzyna Roszkowska, Stephen Holman, Simon Hubbard, Robert Beynon, Malgorzata Adamczyk

## Abstract

Protein biosynthesis is energetically costly, is tightly regulated and is coupled to stress conditions including glucose deprivation. RNA polymerase III (RNAP III) driven transcription of tDNA genes for production of tRNAs is a key element in efficient protein biosynthesis. Here we present an analysis of the effects of altered RNAP III activity on the *Saccharomyces cerevisiae* proteome and metabolism under glucose rich conditions. We show for the first time that RNAP III is tightly coupled to the glycolytic system at the molecular systems level. Decreased RNAP III activity or the absence of the RNAP III negative regulator, Maf1 elicit broad changes in the abundance profiles of enzymes engaged in fundamental metabolism in *S. cerevisiae.* In a mutant compromised in RNAP III activity there is a repartitioning towards amino acids synthesis *de novo* at the expense of glycolytic throughput. Conversely, cells lacking Maf1 protein have greater potential for glycolytic flux.

## Introduction

Regulation of glycolytic flux is a long standing, but still highly relevant, issue in biology and pathobiology. Glycolytic performance is connected to enzymes abundance, cell fermentative activity and proliferation, all hallmarks of the “Warburg effect”. Both *Saccharomyces cerevisiae* and mammalian cells can sense glycolytic state/flux intracellularly, a dominant signal over that of external nutritional status [1–4]. In *Saccharomyces cerevisiae* under favorable growth conditions, high glycolytic activity elicits rapid cell growth, due to robust synthesis of proteins and biomass expansion [5–7]. Nutrient limited growth, on the other hand, is associated with a down regulation of transcription and protein synthesis to reduce demands on the ribosomal machinery and an appropriate supply of amino acids and tRNAs.

As key players in protein synthesis, transfer RNAs are synthesized by RNA Polymerase III (RNAP III), which is also responsible for the transcription of other specific products such as ribosomal 5S rRNA and spliceosomal U6 snRNA. RNAP III activity is regulated by extracellular glucose levels [8,9]. The only known direct regulatory factor of RNAP III in *S. cerevisiae* is the protein Maf1, a mediator of a range of stress signals [10–13] conserved from yeast to human [14]. Yeast Maf1 inhibits RNAP III activity reversibly under carbon source starvation and oxidative stress, reducing tRNA transcript levels [15]. Although the *MAF1* gene is not essential for yeast viability, *maf1Δ* cells are unable to repress RNAP III [15–18].

Under favorable growth conditions, Maf1 is an interaction partner of several cytoplasmic proteins playing different biological functions (Fig 1 A and B), but its function in the cytoplasm is unknown. Maf1 is a target of several kinases and phosphorylation patterns may dictate cellular localization [15, 19–25] (Fig 1 B). Although *MAF1* deletion is not lethal under optimal growth conditions, deletion mutants display high tRNA transcription with diminished growth on non-fermenting carbon sources at 30 °C; it becomes lethal, however, at elevated temperatures. The low growth rate results from a decrease in steady state mRNA levels of *FBP1* and *PCK1* genes encoding the key gluconeogenesis enzymes fructose 1,6 bisphosphatase (Fbp1) and phosphoenolpyruvate carboxykinase (Pck1) [10,26]. Intriguingly, this *maf1Δ* growth defect on non-fermentable carbon sources is suppressed by point mutation (*rpc128-1007*) in the second largest RNAP III subunit *RET1*/C128 [13]. tRNA transcription levels in this *rpc128-1007* mutant are very low, which suggests that the temperature-sensitive lethality of *maf1Δ* can be rescued by attendant reduction of RNAP III activity, or a critical process affected by this transcription. The *maf1Δ* and *rpc128-1007* strains have different phenotypes, not only in growth on non-fermentable carbon sources, but also in preference towards glucose utilization, in excess glucose [13,27]. Transcription of the high affinity glucose transporter genes *HXT6, HXT7* is decreased in *maf1Δ*, but increased over WT in the *maf1Δ* second-site suppressor rpc128-1007 [27], suggesting differences in glucose utilization.

**Fig 1.**
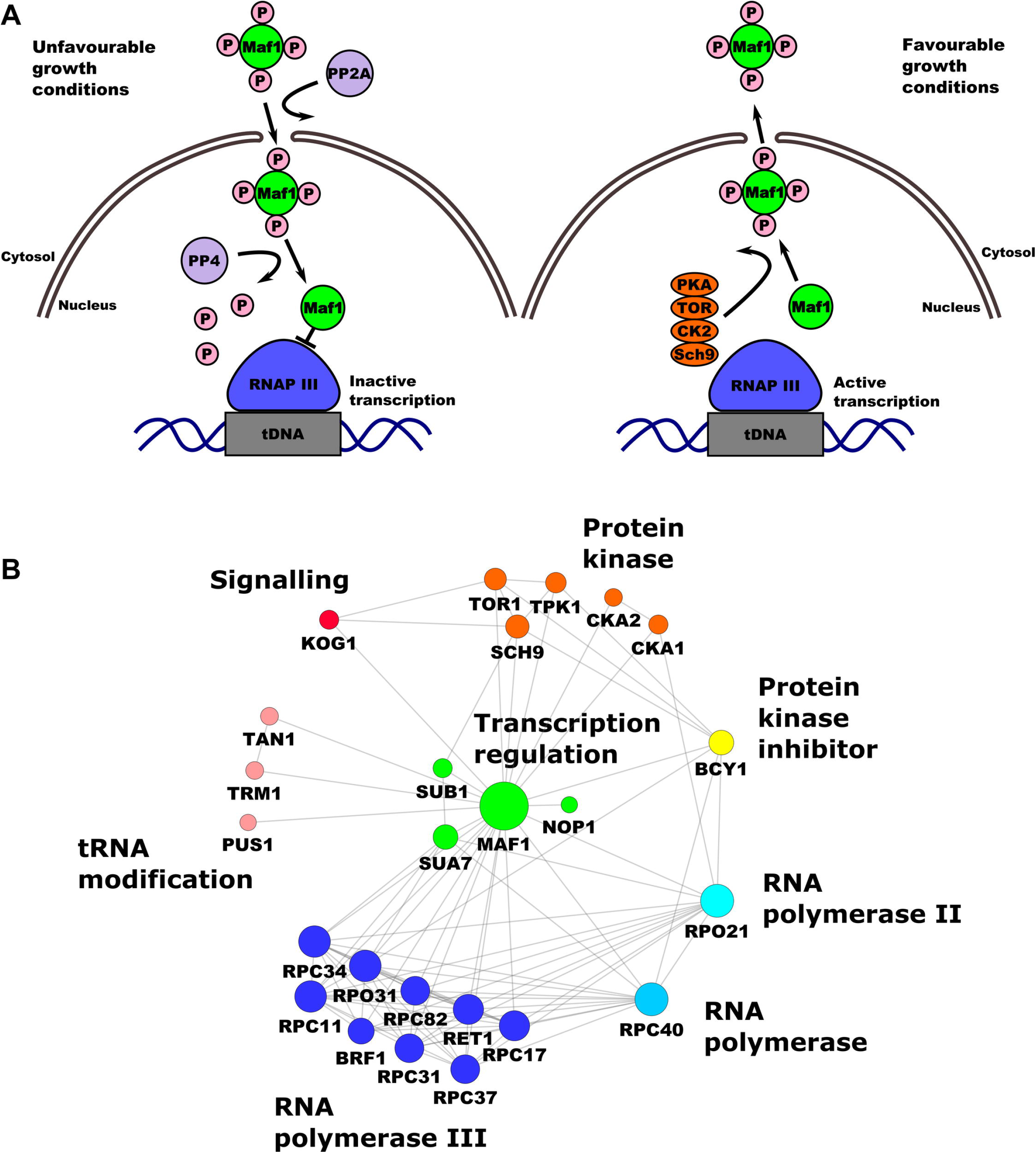
RNAP III regulation by Maf1 and Maf1 interaction network. A) RNAP III transcription repression is regulated by Maf1. Phosphorylation and dephosphorylation events are involved in the mobility and transportation of Maf1 through the nuclear membrane in which a group of protein kinases are involved in the control of Maf1 nuclear localisation responding to stress events. Maf1 produces transcriptional repression on RNAP III by inducing conformational changes. B) Maf1 protein-protein interaction network. Experimental interactions from STRING database are shown. Nodes have been coloured by protein activity in which different protein complexes related to tRNA modification and transportation can be observed. green: transcription regulation; *MAF1*: negative regulator of RNAP III, *SUB1*: *Sub1* transcriptional regulator facilitating elongation through factors that modify RNAP II, role in hyper-osmotic stress response through RANP II and RNAP III, negatively regulates sporulation [121–123], NOP1: Nop1, histone glutamine methyltransferase, modifies H2A at Q105 in nucleolus that regulates transcription from the RNAP I promoter involved in C/D snoRNA 3’end processing. Essential gene leads to reduced levels of pre-rRNA species and defects in small ribosomal subunits biogenesis [124–126], *SUA7*: transcriptional factor TFIIB, a general transcription factor required for transcription initiation and start site selection by RNAP II [127,128] - Sub1 interaction with TFIIB, [129]. Marine blue: RNAP III holoenzyme subunits, red: protein kinases, *KOG1*: Kog1 the component of the TPR complex, Kog1 depletion display the starvation-like phenotypes-cell growth arrest, reduction in protein synthesis, glycogen accumulation, upregulation in the transcription of nitrogen catabolite repressed and retrograde responses genes conserved in from yeast to man is the homolog of the mammalian TORC1 regulatory protein RAPTOR/mKOG1 [82,130], TOR1 mediates cell growth in response to nutrient availability and cellular stress by regulating protein synthesis, ribosome biogenesis, autophagy, transcription activation cell cycle [131,132] yellow: PKA kinase inhibitor protein *BCY1*, pink: tRNA modification *TAN1*: tRNA modifying proteins Tan1 (responsible for tRNA^SER^ turnover [133]), *TRM1*: Trm1 tRNA methyltranspherase produces modified base N2, n2 dimethylguanosine in tRNA in nucleus and mitochondrion [134], *PUS1*:.*PUS1* associated with human disease [135], introduces pseudouridines in tRNA, also as on U2 snRNA and pseudouridylation of some mRNA [136,137].),blue: RPC40 (AC40) is a common subunit to RNAP I and III conserve in all eukaryotes [138,139] light blue: *RPO21*: largest subunit of RNAP II, which produces all nuclear mRNAs, most snoRNAs and snRNA and the telomerase RNA encoded by *TLC1* [140,141], (according to Saccharomyces Genome Database).

We wished to explore the potential for a feedback loop between control of glycolytic flux and RNAP III in yeast cells by label free proteomics, which revealed changes in abundance of a large group of proteins in *maf1Δ* and *rpc128-1007* strains, supported with targeted analysis of specific metabolites. We provide novel molecular data which is able to explain the severe reduction in growth rate caused by RNAP III mutation *rpc128-1007* through cellular processes that facilitate efficient glucose metabolism in the *MAF1* deletion strain on glucose. Changes in protein profiles impact several metabolic pathways, suggesting differences in cellular metabolic homoeostasis in the mutant strains and providing an alternative explanation for *maf1Δ* lethality on non-fermentable carbon sources. Finally, using yeast as a model organism, which is often used for studies of the “Warburg effect”, we established direct metabolic relationship between the capacity of the glycolytic pathway and transcription of non-coding genes, which can explain why several cancerous cell lines exhibit higher RNAP III activity, creating a new perspective on glucose flux modification via manipulation of the RNAP III holoenzyme as an novel therapeutic strategy.

## Materials and Methods

### Yeast strains and media

The following strains were used: wild-type MB159-4D [28] with unchanged RNAP III activity, the MA159-4D *maf1::URA3* [27] *MAF1*-deficient mutant with elevated RNAP III activity and MJ15-9C mutant [13] with a single point mutation in the *RET1/C128* RNAP III subunit with reduced polymerase activity. Yeast strains were cultured in rich medium (YP; 1% yeast extract, 1% peptone) supplemented with either 2% glucose (YPD) or 2% glycerol (YPGly) as a carbon source. Overnight cell cultures were grown in YPD medium. Cells were harvested by centrifugation (2000 rpm, RT) and washed twice with fresh, sterile YPD or YPGly medium. Yeast cells were diluted to A_600_ ≈ 0.1 and grown in YPD or YPGly until exponential phase (A_600_ ≈ 1.0). All yeast cultures were incubated in 30°C with agitation 250 rpm. *GCN4-3HA* DNA construct for chromosomal C-terminus fusion was prepared as described previously [27,29]. Hexokinase isoforms (*HXK1, HXK2, GLK1*) single and double gene deletions were created by transforming haploid yeast strains with appropriate PCR fragments. For *HXK1* deletion amplification of *His3MX6* cassette on *pFA6-VC155-His3MX6* plasmid DNA was done. DNA constructs for obtaining *HXK2* and *GLK1* deficient strains were amplified on gDNA of BY4741 *glk1*∆ and BY4741 *hxk2*∆ (Euroscarf). High-efficiency yeast transformation using LiAc/SS carrier DNA/PEG method was used according to Gietz [30]. All yeast strains are listed in Table 1.

**Table 1.**
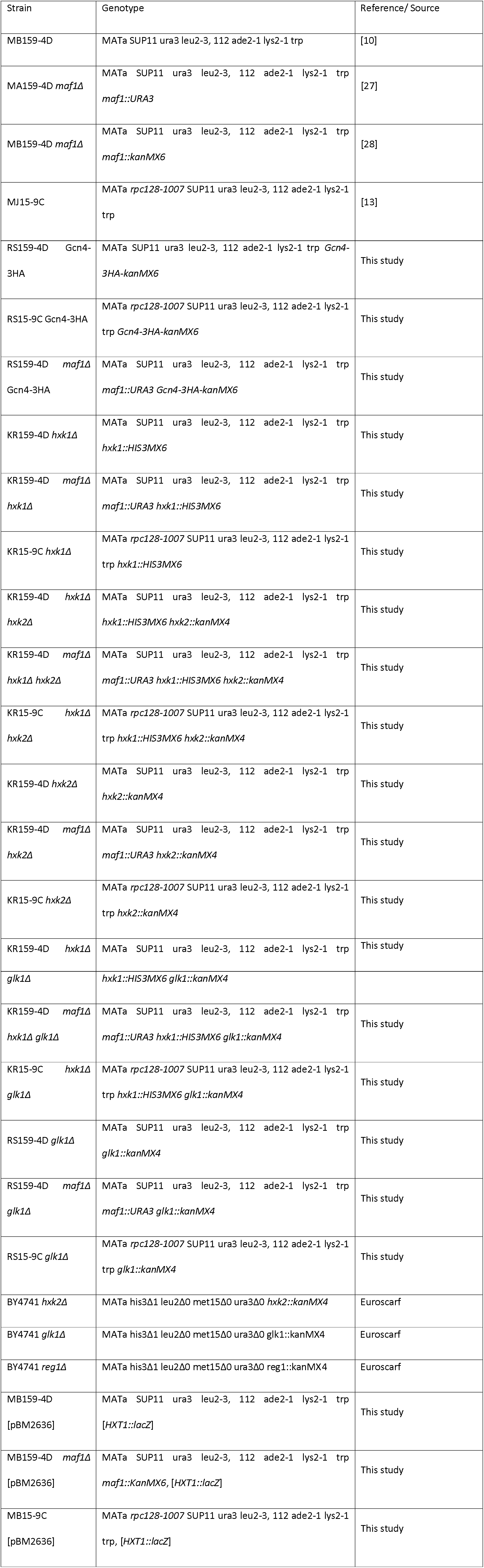
Yeast strains used in the study.

### Proteomic analysis

Samples (approx. 15 ml culture medium) - corresponding to 25 x 10^6^ cells as determined in a cell count on hemocytometer were analysed by global label-free proteomics. Cells were spun down (10 min, 4°C) and the pellets flash frozen in liquid N_2_ for storage. Cells were resuspended in 50 mM NH_4_HCO_3_, protease inhibitor ROCHE mini complete protease inhibitor. The samples were homogenized with MiniBeadbeater 24 (Biospec products) using the 200 μl of glass beads (425-600 μm; Sigma Aldrich) 15 times (3000 hits per min) with a duration of 30 s each with a 1 min cool down period in between each cycles. The cells were further centrifuged (10 min, 13000 rpm, 4°C). 250 μl fresh breaking-buffer was added to pellets and cells were washed by vigorous vortexing. The wash and cell debris were collected as flow through. Each flow through and supernatants from previous steps were combined. Protein concentration was determined with a Bradford assay [31].

A volume equivalent to 25 x 10^6^ cells of each homogenate was removed, diluted with 25 mM AMBIC containing 0.05% Rapigest (Waters, Manchester) and shaken (550 rpm, 10 min, 80°C). The samples were then reduced (addition of 10 μl of 60 mM DTT and incubation at 60°C for 10 min) and alkylated (addition of 10 μl of 180 mM iodoacetamide and incubation at room temperature for 30 min in the dark). Trypsin (Sigma, Poole, UK, proteomics grade) was reconstituted in 50 mM acetic acid to a concentration of 0.2 μg/μl and 10 μl was added to the sample followed by overnight incubation at 37°C. The digestion was terminated and RapiGest™ removed by acidification (1 μl of TFA and incubation at 37°C for 45 min) and centrifugation (15,000 x g, 15 min). To check for complete digestion each sample was analyzed pre- and post-acidification by SDS-PAGE.

For LC-MS/MS analysis, a 2 μl injection of each digest, corresponding to approximately 25 x 10^4^ cells, was analyzed using an Ultimate 3000 RSLC^™^ nano system (Thermo Scientific, Hemel Hempstead) coupled to a QExactive^™^ mass spectrometer (Thermo Scientific). The sample was loaded onto the trapping column (Thermo Scientific, PepMap100, C18, 300 μm X 5 mm), using partial loop injection, for 7 min at a flow rate of 4 μl/min with 0.1% (v/v) FA. The sample was resolved on the analytical column (Easy-Spray C18 75 μm x 500 mm 2 μm column) using a gradient of 97% A (99.9% water: 0.1% formic acid) 3% B (99.9% ACN: 0.1% formic acid) to 60% A: 40% B over 90 min at a flow rate of 300 nL min^-1^. The data-dependent program used for data acquisition consisted of a 70,000 resolution full-scan MS scan (AGC set to 1e^6^ ions with a maximum fill time of 250 ms) the 10 most abundant peaks were selected for MS/MS using a 17,000 resolution scan (AGC set to 5e^4^ ions with a maximum fill time of 250 ms) with an ion selection window of 3 m/z and a normalised collision energy of 30. To avoid repeated selection of peptides for MS/MS the program used a 30 s dynamic exclusion window.

### Label-free quantification

The raw data from the mass spectrometer was then processed using MaxQuant (MQ) software version 1.5.3.30 [32]. Protein identification was performed with the built-in Andromeda search engine, searching MS/MS spectra vs. the *S. cerevisiae* strain ATCC 204508/S288c downloaded from UniProt. (https://www.uniprot.org/proteomes/UP000002311). The following parameters were used; digest reagent: trypsin, maximum missed cleavages: 2, modifications: protein N-terminal acetylation and methionine oxidation, with a maximum of five modifications per peptide. The false discovery rate (FDR) for accepted peptide spectrum matches and protein matches was set to 1%. For protein quantification, the ‘match between runs’ options was selected. Label free quantification was performed with the MaxLFQ algorithm within MaxQuant (MQ), based on razor and unique peptides. All other MQ parameters were left at default values.

### Protein significance testing

To determine statistically significantly changing proteins with respect to the wild-type strain we used the MSstats package [33] in the R environment. Protein identities, conditions, biological replicates and intensities were directly uploaded from the MaxQuant output. Protein ID information was obtained from the ‘proteinGroups.txt’ file, conditions and biological replicates from the ‘annotation.csv’ file, and intensities from the ‘evidence.txt’ file. Data normalization was performed using the ‘equalizeMedians’ option and summarization using the Tukey’s median polish option. Following this, a condition comparison was performed using the ‘groupComparison’ option from where the log_2_ fold changes and adjusted *p*-values were obtained.

### Functional analysis

Gene ontology enrichment analysis was performed with the online application Panther (34), directly on the Gene Ontology Consortium webpage (http://pantherdb.org/). The background set consisted of all proteins identified in a given MS experiment. Protein changes were mapped to central carbon and amino acid metabolic pathways following KEGG database [35] guidelines. Maf1 protein-protein interactions (PPI’s) were obtained from the STRING [36] database and clustered with the Cytoscape [37] tool.

### Transcription Factor target enrichment analysis

For transcription factor (TF) target enrichment analysis, all proteins with an adjusted p value below 0.05 from both comparisons (WT - *rpc128-1007* and WT - *maf1Δ*) were uploaded to the GeneCodis tool [38]. Proteins were classified according to their positive or negative fold change and the background set consisted of all proteins identified in the given MS experiment. All statistical parameters were left as default. Adjusted *p*-values were obtained indicating those statistically significantly TFs being active according to their known target proteins.

### Western blotting

The total cellular proteins from Gcn4-3HA expressing yeast cells were extracted as described previously [27]. Protein extracts were separated by 12% SDS-PAGE and transferred to nitrocellulose membrane by electrotransfer (1 h, 400 mM, 4°C). For detection of HA-tagged proteins, monoclonal mouse anti-HA (1:3330, Sigma, H3663) and polyclonal goat anti-mouse antibodies (1:2000, Dako P0447) conjugated with horseradish peroxidase (HRP) were used. For Vma2 protein detection, monoclonal mouse anti-Vma2 antibodies (1:4000, Life Technologies, A6427) were used.

### RNA isolation and RealTime PCR quantification

RNA isolation and real-time PCR amplification was performed as described previously [27]. Isolated RNAs were examined by SYBR GREEN-based Real-time PCR. Oligonucleotide sequences of the primers used in Real-time PCR experiment for GCN4 were taken from Cankorur-Cetinkaya *et al*. [39]. Samples were normalized to two reference genes - *U2* spliceosomal RNA (U2) and small cytosolic RNA (*SCR1*). Expression levels in WT strain (MB159-4D) was taken as 1.0. The relative expression (mean±SD) were calculated for at least three independent biological replicates. Statistical significance of *p*-values were determined by t-student test.

### Enzymatic assays

All yeast strains including transformants carrying *pBM2636* plasmid [40] for measurement of β-galactosidase activity were cultivated in rich medium supplemented with 2% glucose (YPD) or 2% glycerol (YPGly) at 30°C with agitation of 250 rpm until reached A_600_ ≈ 1.0. Yeast cultures were harvested at 5000 rpm at 4°C and washed twice with 10 mM potassium phosphate buffer, pH = 7.5. Cells for Hxk, Pgk1, Cdc19 and Zwf1 activity assays were suspended in 100 mM KPi pH = 7.5, for β-galactosidase in 50 mM potassium phosphate buffer pH = 7.0, rapidly frozen in liquid nitrogen and stored at −20°C. Samples were washed twice with sonication buffer (100 mM potassium phosphate buffer, pH = 7.5, 2 mM MgCl_2_) or 50 mM potassium phosphate buffer pH = 7.0 in the second case and disintegrated with Mini-Beadbeater 24 (Biospec products) using glass beads (425– 600 μm; Sigma Aldrich). Hexokinase (EC 2.7.1.1) activity was measured according to Adamczyk [41], phosphoglycerate kinase (Pgk1, EC 2.7.2.3) according to De Winde *et al* [42], pyruvate kinase (Cdc19; EC 2.7.1.40) according to Gruning *et al* [43], glucose-6-phosphate dehydrogenase (Zwf1, EC 1.1.1.49) according to Postma *et al*. [44], β-galactosidase according to Smale 2010 (45), and catalase (EC 1.11.1.6) according to Beers and Sizer [46]. All assays were performed for at least three independent biological replicates.

### Glycogen, trehalose and fructose 1,6 bisphosphate measurement

The glycogen and trehalose content was measured in yeast cells, grown in YPD until A_600_ ≈ 1.0. Cell preparation and extraction was as described in Rossouw *et al*. 2013 [47]. Glycogen determination was as described by Parrou and Francois [48]. Glucose concentration from glycogen enzymatic breakdown was determined by the glucose (HK) Assay Kit according to the manufacturer’s protocol (Sigma Aldrich, GAHK-20). Trehalose content was measured using Trehalose Assay Kit (Megazyme International Ireland, Wicklow, Ireland) according to manufacturer’s protocol. Fructose 1,6 bisphosphate was measured according to Peeters *et al*. (2017) [4] with minor modifications.

### Determination of yeast fermentative capacity

Fermentative capacity assays were performed as described by van Hoek *et al*. [49] with minor changes. The fermentative capacity can be defined as the specific maximal production rate of ethanol per gram of biomass (mmol/g/h) under anaerobic conditions at excess of glucose. Samples corresponding to 60-70 mg dry weight were harvested by centrifugation at 5000 rpm at 4°C. Cells were washed twice with synthetic medium CBS-without carbon source and resuspended in CBS (-C) to make 2% wet weight suspensions. Analysis was performed in a thermostatted (30 °C) vessel. Cells were flushed with N_2_ gas at a flow rate of approximately 0.6 L/h and glucose was added to a final concentration of 10 g/liter. Samples for measurement of ethanol were collected every 5 min incubated with 35% (w/v) perchloric acid on ice for 10 min and neutralized with KOH before centrifugation at 13000 rpm and stored in −20°C freezer. The ethanol production of each strain was normalized to the dry weight of the culture. Ethanol and glycerol in supernatants were determined with enzymatic assays according to manufacturer (Megazyme).

### Biomass determination

Sample suspensions of 1 ml volume in duplicates were filtered over pre-weighted nitrocellulose filters (pore size, 0.45 μm; HAWP04700). After removal of medium, the filters were washed with demineralized water, dried in an oven overnight and weighted [49].

## Results

### Overall proteome profiling and changes

We hypothesized that perturbations in RNAP III activity would impact on global expression of the proteome. The lack of the negative regulator of RNAP III, Maf1, as well as the loss of function due to a point mutation in the RNAP III *RET1*/C128 subunit, would be expected to elicit broad changes in the *Saccharomyces cerevisiae* proteome. We aimed to identify proteins, the changed abundance of which, could explain the diminished growth of *rpc128-1007* on glucose and de-repression of *HXT2* and *HXT6/7* genes that encode high affinity glucose transporters, when compared to *maf1Δ* under glucose rich conditions [27]. To examine the phenomenon we performed a systematic comparative analysis of *maf1Δ* and *rpc128-1007* mutants using label-free proteomics.

Reproducible, deep proteome coverage was obtained (Fig 2 A) for *maf1Δ* and *rpc128-1007* mutants grown under glucose rich conditions. The proteomics data were of high quality, with replicates clustered together and no systematic difference reflecting sample preparation bias (Fig 2). In total, over 2,300 protein groups were identified and quantified. As anticipated, there was considerable overlap between the *maf1Δ* and *rpc128-1007* proteomes (Fig 3). Differential proteome analysis was carried out pairwise in two sets as follows: WT vs *rpc128- 1007* mutant and WT vs *maf1Δ* mutant, resulting in 2,294 quantified proteins common to all strains. A subset of statistically significant changes revealed 249 proteins that were common to both comparisons (with an adjusted *p*-value < 0.05). This subset of 249 proteins were clustered into coherent groups (see Material and Methods) which display common Gene Ontology (GO) annotation, consistent with coordinated regulation of relevant biological processes (Fig 3 B). Some of the groups exhibited parallel changes in the two strains (Groups 1-4) whereas others highlighted divergent, essentially reciprocal, functions in the two strain (i.e. Groups 5–6). These unbiased clusters show enrichments for concerted biological functions, embodied by the limited subset of GO term enrichments listed in Fig 3B, including elements of amino acid and monosaccharide/carbohydrate metabolism. In both strains, enzymes of gluconeogenesis and the glyoxylate cycle were *decreased* when compared to the reference strain grown under the same glucose repression conditions (Group 2). Groups 7 and 9 also show decreased protein abundance with respect to wild type, and are similarly enriched in enzymes from the TCA cycle, purine ribonucleotide biosynthetic pathways, *de novo* inosine monophosphate biosynthesis, and also mitochondrial transmembrane transport. In contrast, many of the enzymes involved in *de novo* amino acid synthesis were increased in abundance in both *maf1Δ* and *rpc128-1007* (Groups 1, 3, 4). The negatively correlated, reciprocally altered groups (Groups 5, 6 and 8) were consistent with shifts in trehalose biosynthesis, pentose phosphate pathway (PPP) activity, oxidative stress, oxidation-reduction processes, glycine catabolic processes, replicative cell aging and alcohol production.

**Fig 2.**
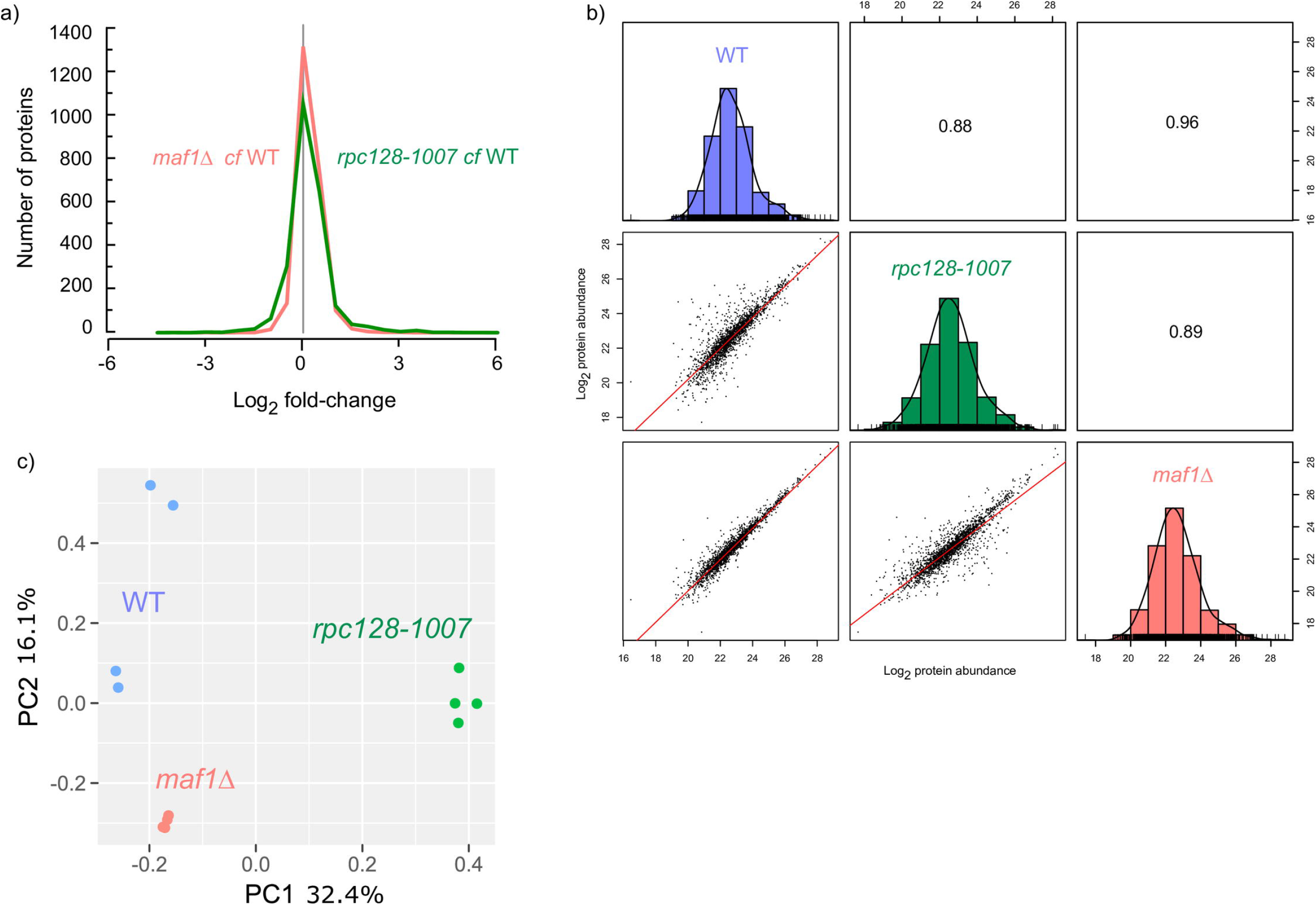
Proteome signature of *maf1Δ* and *rpc128-1007* mutants compared to wild-type strain. A) Histogram of proteins present on both mutants organized according to their corresponding Log_2_ fold change expression. B) Comparative scatter plots and histograms of the different strains. The Log_2_ transformed protein abundances of proteins present in the WT, *rpc128-1007* and *maf1 Δ* strains are plotted against one another along with their distribution. The number shown is the Pearson correlation coefficient between the two relevant strains. C) Principal component analysis (PCA) based on proteins present on all biological replicates.

**Fig 3.**
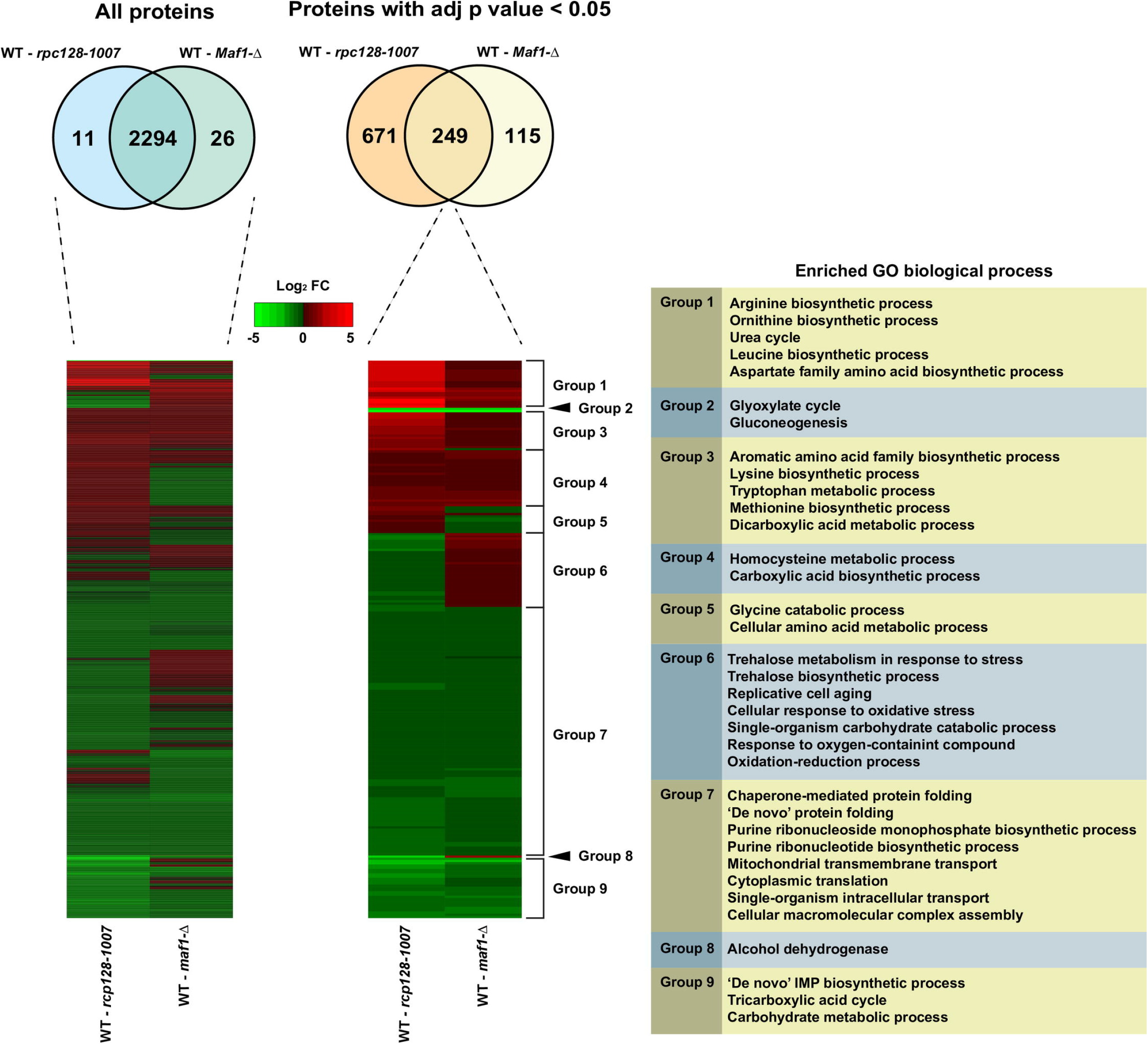
Increased and decreased protein abundance is presented relative to the wild-type strain for both *maf1Δ* and *rpc128-1007* mutants. All those statically significant proteins with an adjusted *p*-value < 0.05 overlapped between both comparison were then subjected to a hierarchical clustering. This clustering analysis created different groups showing the similarities and differences between both mutants with clusters enriching to biological processes related to amino acid and carbohydrate metabolism, response to stress, and respiratory processes.

### Key enzymes of the glyoxylate cycle are reduced in abundance in both *maf1Δ* and *rpc128- 1007* mutants

In both strains phosphoenolpyruvate carboxykinase (Pck1) and malate synthase 1 (Mls1) were reduced (Fig 4). These enzymes direct acetyl-CoA to malate and oxaloacetate that in turn can be metabolized to phosphoenolpyruvate for gluconeogenesis. The enzymes are components of the glyoxylate cycle that allows yeast cells to metabolize non-fermentable carbon sources, including fatty acids. The mechanism governing glucose repressed genes is particularly important in the RNAP III compromised mutant as well as in Maf1 deprived cells due to the previously reported growth perturbations of *maf1Δ* on non-fermentable carbon source. 2-fold decrease in *maf1Δ PCK1* mRNA was reported [26], though under inducing conditions on glycerol. Notably, the relative decrease in Pck1 abundance in Maf1 deficient cells is the largest in our proteomic dataset. The very much decreased Pck1 abundance proves that the enzyme is subject to degradation in a glucose-dependent manner [50] and the mechanism is not perturbed in both the mutants. Other enzymes of the glyoxylate cycle, are concomitantly reduced in *maf1Δ* cells, including malate dehydrogenases Mdh2, Mdh3, isocitrate lyase (Icl1) and glyoxylate aminotransferase (Agx1), the last implicated in glycine synthesis from glyoxylate. In *rpc128-1007*, most of the glyoxylate enzymes as well as those of the TCA cycle were also decreased. Enzymes of the TCA and glyoxylate cycles undergo coordinated transcriptional down regulation [50,51] induced by glucose through the master kinase Snf1/AMPK [52,53] therefore strongly suggesting unperturbed functioning of Snf1 signaling on glucose. In contrast, the other key enzyme of gluconeogenesis, Fbp1 (fructose 1,6 bisphosphatase) that bypasses the physiologically irreversible step in the glycolytic pathway, was increased under glucose deprivation in maf1*Δ*, but was unchanged in *rpc128-1007* mutant.

**Fig 4.**
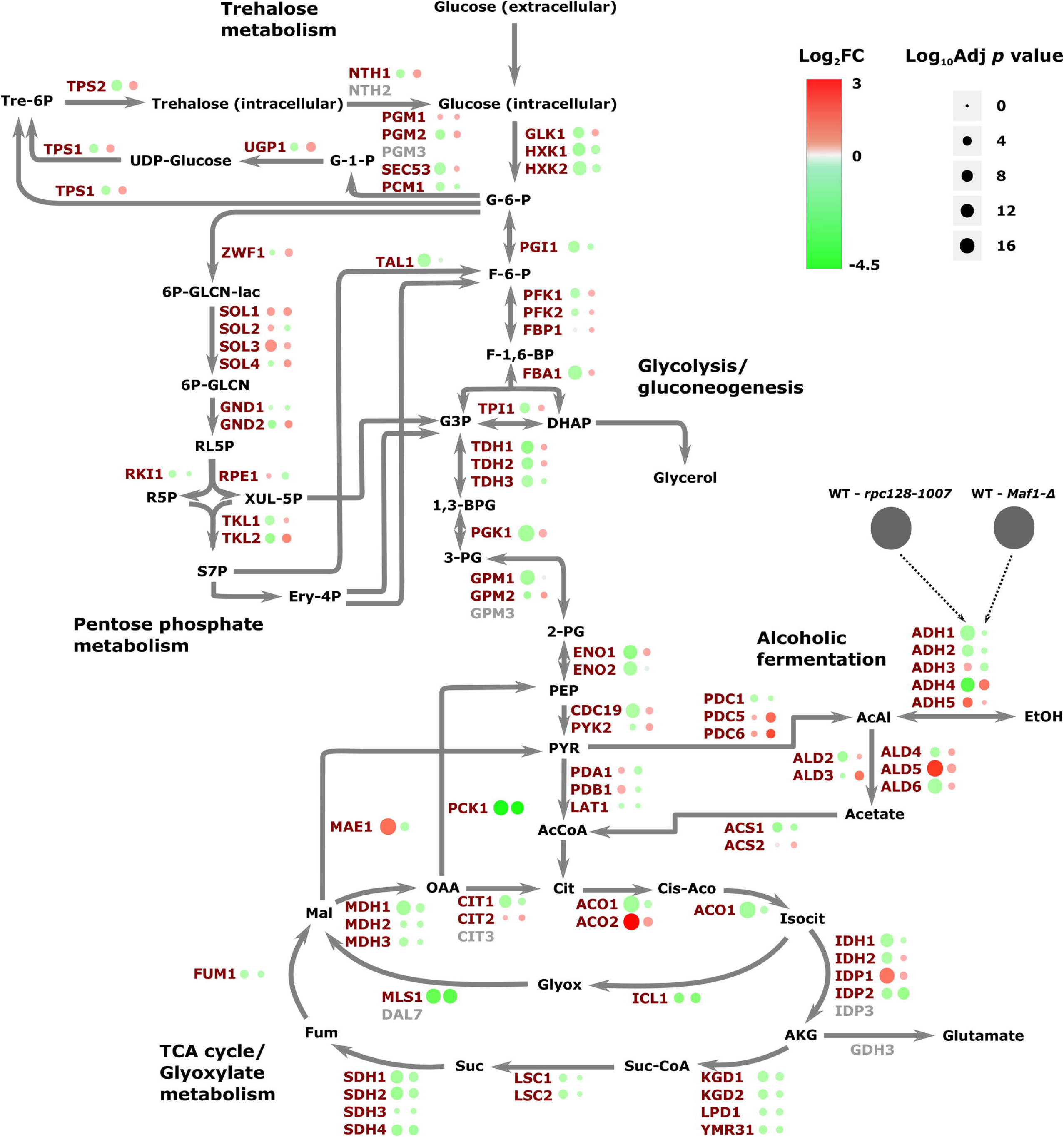
Comparative proteomic profiling of *maf1Δ* and *rpc128-1007* mutants when compared to wild-type strain. The differences in protein abundances are presented on a schematic representation of the central carbon metabolism. Those proteins with and increased abundance are presented in red and those with an decreased abundance in green.

### Reduction in glycolytic enzymes in a RNAP III compromised strain correlates with lower activity of the glucose transporter Hxt1

Proteome analysis captured changes in relative cellular abundances of all glycolytic enzymes and implied a reduced capacity for glycolysis in *rpc128-1007* cells, but an unchanged glycolytic capacity in *maf1Δ* cells (Fig 4). In *rpc128-1007*, enzymes that were significantly decreased (between 2- and 2.6-fold) included glyceraldehyde-3- phosphate dehydrogenase isozyme 1 (Tdh1), enolase (Eno1), glyceraldehyde-3-phophate dehydrogenase isozyme 2 (Tdh2), 3-phosphoglycerate kinase (Pgk1) and glucokinase (Glk1). The lower abundance of the entire complement of glycolytic enzymes is consistent with reduced glycolytic performance in *rpc128-1007*.

Glycolytic flux in *S. cerevisiae* can regulate glucose uptake, at least in part through the activity of glucose uptake mechanisms [54] and in particular, induction and increases of membrane internalization of low-affinity glucose transporters [40, 55–58]. The principal example is Hxt1, only activated when yeast grow in glucose rich media [40]. We explored the potential for changes in glucose transport by measurement of transcriptional derepression of the key gene encoding the major low affinity, high capacity glucose transporter *HXT1*, in *maf1Δ* and *rpc128-1007* using a *HXT1-lacZ* reporter plasmid (Fig 5 B). The *HXT1* gene was strongly activated in *maf1Δ* under glucose rich conditions, whereas a 3-fold lower activity of the *HXT1* promoter was observed in *rpc128-1007* cells in the same conditions. When assessed in cells grown on glycerol, expression of *HXT1–lacZ* reporter was decreased in all strains (Fig 5 B) as expected [40]. Consistent with the reports on mutants in genes of the glycolytic pathway, which are blocked in glycolysis [59], the *HXT1* expression was reduced in RNAP III compromised yeast suggesting that in this mutant, supply of glucose for a functional glycolytic pathway cannot be maintained.

**Fig 5.**
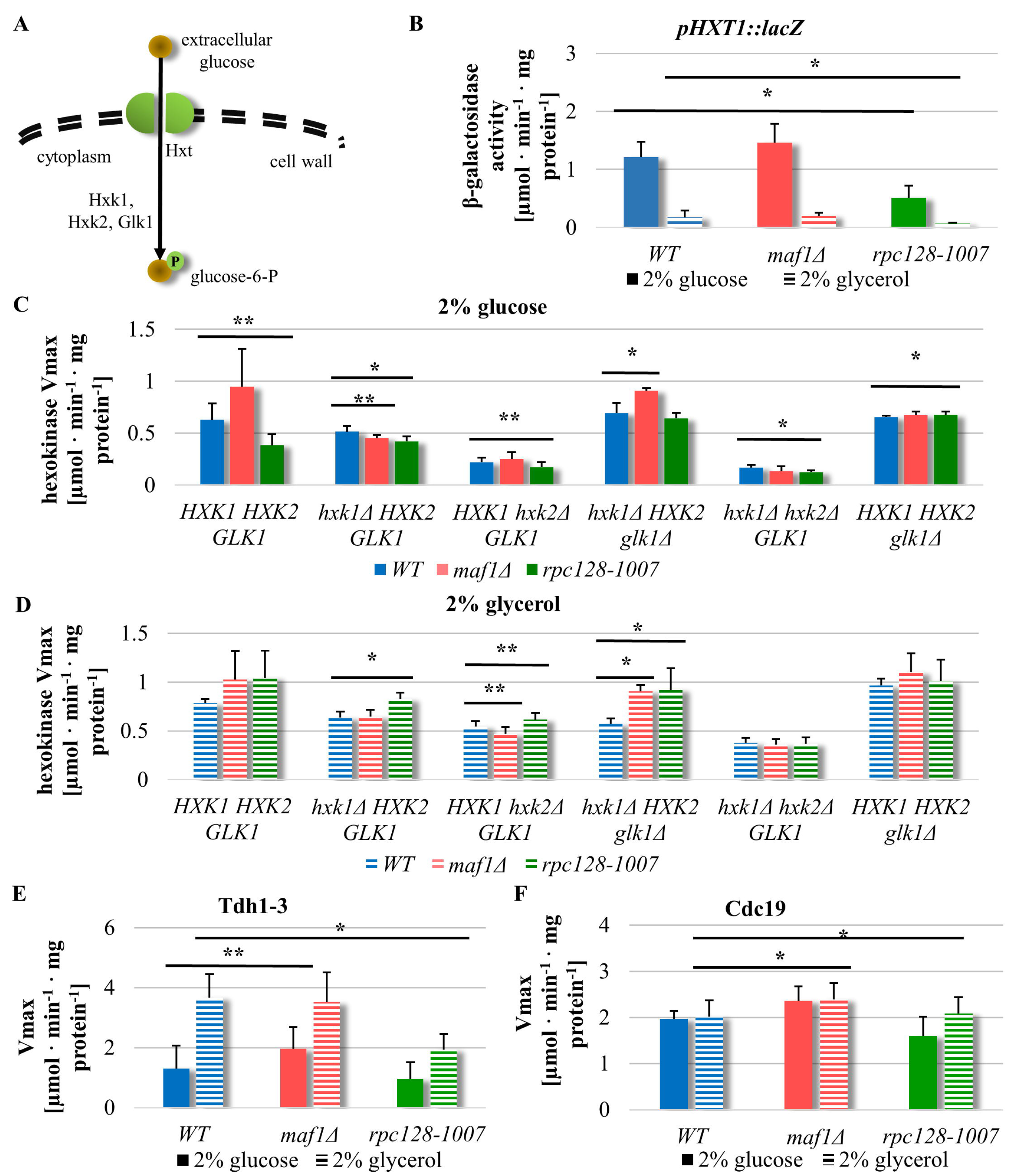
Opposite effects have been observed in *HXT1* promoter activity in strains with altered RNAP III. (A) Schematic representation of glucose uptake and phosphorylation in yeast cells. Hxt – hexose transporter, P – phosphorylation, Hxk1 – hexokinase 1, Hxk2 – hexokinase 2, Glk1 – glucokinase 1. WT, *maf1*Δ, *rpc128-1007* yeast cells and single or double *HXK1, HXK2, GLK1* knockouts strains in WT, *maf1*Δ and *rpc128-1007* genetic background were cultured in YPD (C) or YPGly (D) rich medium under either inducing (2% glucose) or repressing (2% glycerol) conditions. Maf1 deficiency increases *HXT1* expression (B) on glucose and Hxk2 activity regardless carbon source (C, D). Metabolic effects observed in *rpc128-1007* correlate with decreased *HXT1* expression (B) and decreased hexokinase activity in glucose rich medium (C), but increased hexokinase activity in glycerol rich medium (D). Compromised RNAP III and *maf1*Δ have effect on enzymes in lower glycolysis: Tdh and Cdc19 activities (E and F). The WT strain (MB159-4D), *maf1*Δ and *rpc128-1007* mutant strains were grown under 2% glucose and 2% glycerol conditions. The experiment was performed in cell-free extracts isolated from the aforementioned strains. Data are expressed as the mean obtained from at least three independent experiments measured in triplicate. The + standard deviations are shown. Enzymatic assays were performed in cell-free extracts. The reaction rates were monitored by measuring NADH concentration change over time at 340 nm. V_max_ mean value is expressed as μmol·min^-1^·mg^-1^ protein (C, D, E, F). (B) *HXT1* expression was measured in WT [pBM2636], *maf1*Δ [pBM2636] and *rpc128-1007* [pBM2636] strains by using the *lacZ* reporter gene system [40]. β-galactosidase activity was assayed in cell-free extracts. The error bars indicate the standard deviation from three independent transformants assayed in triplicate. Asterix (*) indicate *p*-value < 0.05 and double asterix (**) illustrate *p*-values < 0.1 according to t-student test.

### Positive relationship between Hxk2, Tdh and Cdc19 enzyme activities and the potency of RNAP III dependent transcription

Since *HXT1* gene expression was elevated in *maf1Δ*, but decreased in *rpc128-1007*, we measured the activity of selected glycolytic enzymes *in vitro* [41] in both mutant strains and evaluated the relationship between activity changes and changes in protein abundance assessed by proteomics. *S. cerevisiae* encodes three isoenzymes with hexokinase activity (Fig 5 A). Proteomic analysis suggested that Glk1 was the isoenzyme phosphorylating glucose in *maf1Δ* (Fig 4), with increased abundance whereas Hxk1 and Hxk2 were observed decreased in this strain. For *rpc128-1007*, the protein abundance of all enzymes conferring hexokinase activity was decreased (Fig 4). We therefore grew the three strains in rich media supplemented with 2% (w/v) glucose and measured the hexokinase reaction (V_max_) in cell free extracts. Total hexokinase activity was increased in *maf1Δ* and reduced in *rpc128-1007* (Fig 5 C, solid bars).

To quantify the activities of the individual hexokinase enzymes we designed and constructed deletion mutants of hexokinases in the three strains (Fig 5 C, D). Quantification of glucose phosphorylation activity in single and double null mutants of genes encoding hexokinases clarifies that hexokinase 2 (Hxk2) is the predominant isoenzyme engaged in glucose phosphorylation in *maf1*Δ. The triple deletion *maf1*Δ *hxk1*Δ *glk1*Δ, in which the only isoform left intact is Hxk2, results in comparable hexokinase activity to the observed in *maf1*Δ deletion strain with all the isoforms present (Fig 5 C). In contrast, the mutants in whom we observe the reverse trend in the enzymatic activity, are the *maf1*Δ *hxk2*Δ double mutant and *maf1*Δ *hxk1*Δ *hxk2*Δ triple mutant. An increase in glycolytic flux is possible to achieve in cells lacking Maf1 despite a decrease in Hxk2 abundance and only a slight increase in Glk1 cellular concentration. Under growth on glycerol, there was an increased contribution of Hxk1 to total hexokinase activity in wild-type, *maf1*Δ and *rpc128-1007* (Fig 5 D), suggesting that the compensation regulatory mechanisms are not perturbed in the two mutant strains. *HXK1* induction by non-fermentable carbon source has previously been reported [60]. Interestingly, on glycerol growth, the total hexokinase activity in *rpc128-1007* was higher than in *maf1*Δ.

Since glyceraldehyde-3-phoshate dehydrogenase (Tdh) and pyruvate kinase (Cdc19) are important providers of NADH and ATP respectively, these enzymes were also assayed. Measuring the activity of controlling and rate-limiting glycolytic enzymes is one of the techniques to estimate carbon flux thought the entire pathway. In yeast, glyceraldehyde-3-phoshate dehydrogenase, placed between upper and lower segments of glycolysis, is considered a rate controlling step of glycolysis [61,62] whereas Cdc19 kinase levels affect the rate of carbon flux and its direction towards pyruvate (PYR) or phosphoenolpyruvate (PEP) under fermentative conditions. The activity of Cdc19 is sufficient to cause a shift from fermentative to oxidative metabolism in *S. cerevisiae* [43,63] and controls glycolytic rate during growth on glucose [64].

In *maf1Δ*, in which there was a small increase of Tdh1, 2 abundance (Tdh1; 0.21 log_2_FC and 0.3 adj. *p* value, Tdh2: 0.16 log_2_FC and 0.37 adj. *p* value), *in vitro* activity was 2-fold higher (Fig 5 E). From proteomics, Tdh activity was slightly lower in *rpc128-1007* cells under the same growth conditions, but Tdh activity measured in *rpc128-1007* grown on glycerol was significantly lower when compared to the reference strain, which suggests that the catalytic activity of the enzyme decreases *in vivo* while the enzyme converts 1,3-bisphosphoglycerate (1,3-BPG) into glyceraldehydes-3-phosphate (G3P) in the reverse direction to glycolysis, when the enzymes is involved in gluconeogenesis in the presence of non-fermentable carbon sources in the medium. This shuttle between the cytosol and the nucleus linking metabolic redox status to gene transcription [65] and contributes to tRNA transport [66].

We further measured activity of the final enzyme in the glycolytic pathway, pyruvate kinase (Cdc19) activity. Cdc19 concentration is comparable in *maf1Δ* and its parental strain. We found (Fig 5 F), that Cdc19 shows significantly lower enzymatic activity in *rpc128-1007* compared to the reference strain grown on glucose, but a slightly elevated activity in Maf1 deficient cells both on glucose and glycerol. Overall, in *maf1Δ* the glycolytic enzymes show higher activity than originally thought judging by proteomics data, whereas Hxk2, Tdh and Cdc19 protein decreased abundance in *rpc128-1007* is fully in agreement with their reduced enzymatic activity. This lead us to the conclusion that glycolytic flux is diminished in *rpc128-1007*, whereas in *maf1Δ* it is not only higher than in *rpc128-1007* but also in WT. Due to the fact that F16BP mediated allosteric control [67–69], but not Cdc19 abundance or phosphorylation, was reported as having a predominant role in regulating the metabolic flux through the pyruvate kinase Cdc19 [70]. Therefore, we measured F16BP intracellular concentration.

### Fructose 1,6 bisphosphate intracellular concentration does not reflect differences in RNAP III activity and glycolytic flux in the mutant strains

The glycolytic metabolite F16BP is a molecule triggering of the metabolic switch from respiration to fermentation in unicellular and higher organisms [3,4,64]. We reasoned that lower glycolytic flux in *rpc128- 1007*, would result in lower fructose 1,6 bisphosphate (F16BP) concentration in this mutant. We measured F16BP in *rpc128-1007* grown under anaerobic conditions. After addition of 100 mM glucose, F16BP concentration sharply increased during the first two minutes to the physiological level observed in the wild-type cells but then declines to a new steady state. In *maf1Δ* cells, the concentration of F16BP is similar to *rpc128-1007* but there is no initial overshoot (Fig 6). The intracellular level of F16BP is therefore unlikely to be the trigger for the perturbed metabolic switching in cells with different RNAP III activity. Another product of glycolysis (or its side branches) may control the transcriptional reprogramming in yeast, particularly when there are multiple-metabolite-responsive elements present at promoters to sense diverse metabolic signals. Additionally, F16BP concentration may affect Cdc19 activity *in vivo*, thus the flux direction.

**Fig 6.**
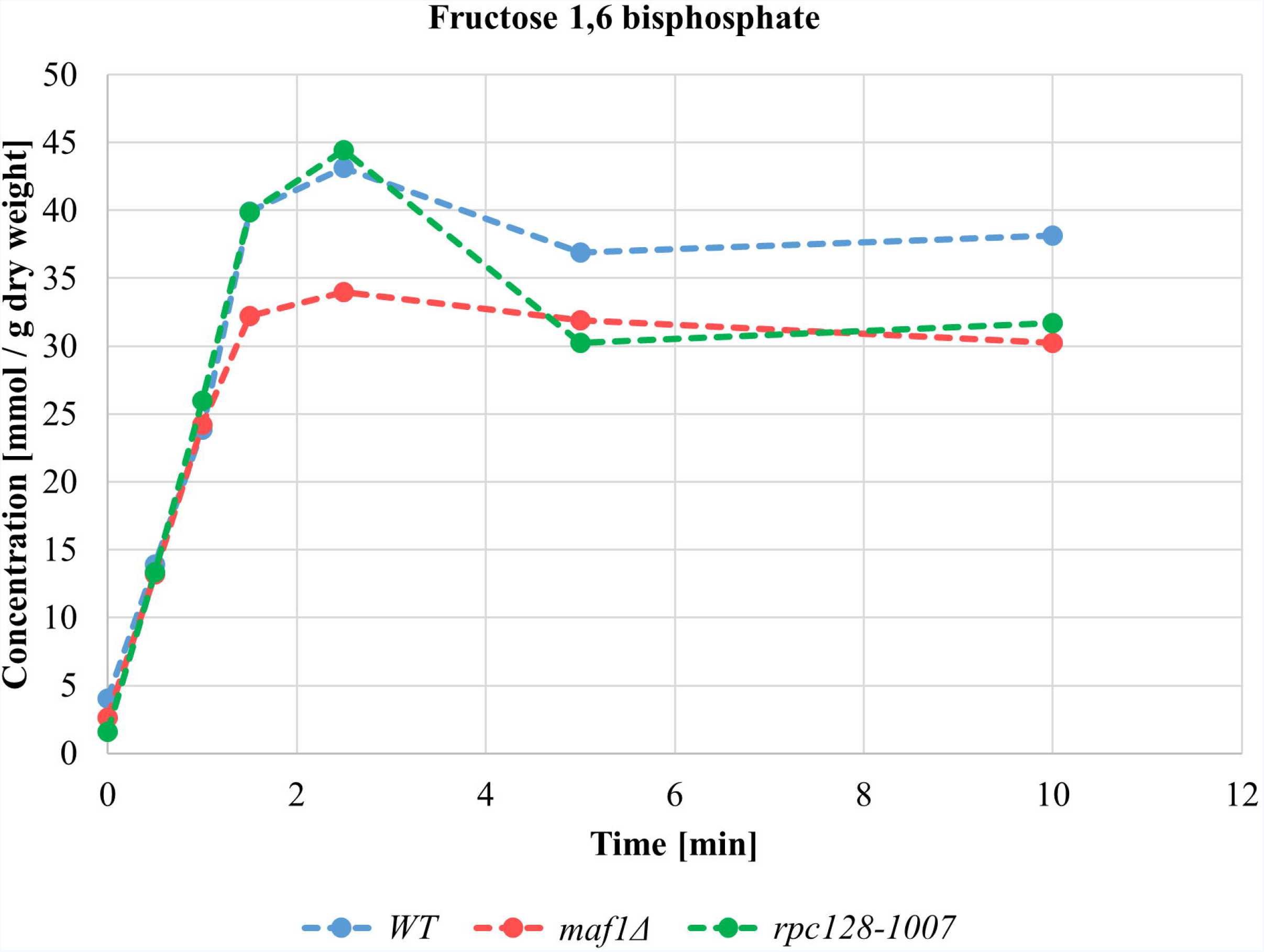
Changes in intracellular concentration of Fructose 1,6 bisphosphate (F16BP). Intracellular fructose 1,6 bisphosphate concentration is lowered in cells with changed RNAP III activity under glucose pulse experiment. Cells were grown in YPD until reaching A_600_ ≈ 1.0, collected washed in minimal medium lacking carbon source (CBS-C) and resuspended in CBS (-C). Analysis was performed in a thermostatted vessel at 30°C. Cells were flushed with Ar_2_gas and glucose was added to a final concentration of 2%. Cell samples suspension were collected in time. Fructose 1,6 bisphosphate content was measured by enzymatic breakdown of NADH monitored by changed absorbance at 340 nm in time according to [4]. Fructose 1,6BP concentration was calculated from a standard curve and standardized to cells dry weight expressed in g. Results are shown as mean value for four biological replicates.

### Higher glucose flux in Maf1 deficient cells results in activation of glycogen and trehalose shunts

We explored the direction of carbon flux in *maf1Δ* in the absence of any increase in F16BP concentration. Yeast cells are equipped to counteract excessive influx of glucose by diversion of glucose into glycogen and trehalose [71]. During exponential growth, glycogen and trehalose biosynthesis play additional roles as part of an adaptive response facilitating survival when the cell is challenged with increased glycolytic flux as a consequence of glucose overflux into a cell. The glycogen shunt prevents accumulation of glycolytic intermediates, particularly F16BP and ATP that otherwise would ultimately lead to perturbation in cell metabolic homeostasis [72–75].

The proteomics data confirmed a strong, negative correlation between RNAP III activity and enzyme abundance in the trehalose and glycogen synthesis pathways, which share common enzymes. UDP-glucose pyrophosphorylase (Ugp1), trehalose-6-P synthases (Tps1 and Tps2) and glycogen synthase (Gsy2) increase in *maf1Δ*, whilst the same proteins were markedly reduced in *rpc128-1007* cells (Fig 3, group 6, Fig 4). The product of Tps1 activity, trehalose-6-phosphate, controls glycolysis by restricting the flow of glucose into the pathway and is an allosteric inhibitor of hexokinase 2 activity [72,76]. We assessed the metabolic allocation of glucose through quantification of trehalose and glycogen content during exponential growth. Both metabolites were 2.5-fold higher in *maf1*Δ in the presence of high glucose. By contrast, *rpc128-1007* cells could accumulate neither glycogen nor trehalose (Fig 7 A and B).

**Fig 7.**
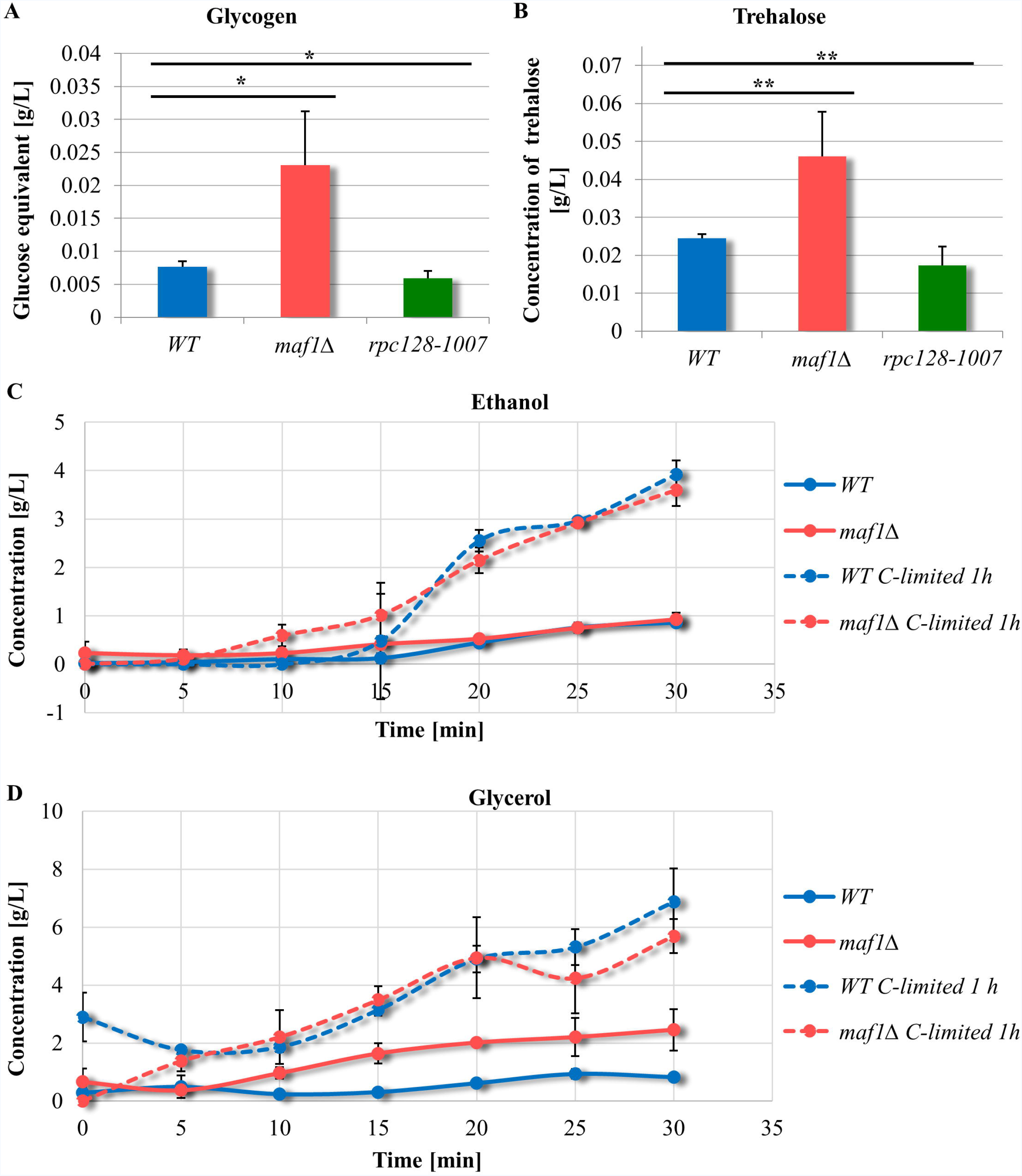
Maf1 deficient yeast strain accumulates glycogen (A) and trehalose (B) during exponential phase. Yeast were cultivated in rich medium supplemented with 2% glucose and harvested by centrifugation at A_600_ ≈ 1.0. Glucose concentration from enzymatic breakdown of glycogen (A) by amyloglucosidase from *A. niger*, was determined by Glucose (HK) Assay Kit (GAHK-20, Sigma). Trehalose (B) content determination assay was performed using Trehalose Assay Kit (Megazyme) according to manufacturer’s protocol. Trehalose and glycogen content is presentment as mean value of at least three independent biological replicates with standard deviations. There were no significant changes in ethanol production rate between wild-type (MB159- 4D) and *maf1Δ* strain (C). *maf1Δ* accumulated glycerol (D). Ethanol and glycerol concentration was determined under Fermentative capacity assay (FCA) conditions in *maf1*Δ strain (C and D). Fermentative capacity assay was performed as described by van Hoek *et al*. (1998) [49] with modifications (for details see Method section). All assays were performed in triplicates. Results are shown as mean concentration [g/L] value with standard deviation in time [min]. ‘C-limited’ stands for ‘carbon-limited conditions’. Asterix (*) indicate *p*-value < 0.05 and double asterix (**) illustrate *p*-values < 0.1 according to t-student test.

In summary, the results are consistent with increased glycolytic flux in *maf1*Δ and conversely, a diminished flux in *rpc128-1007*. Metabolic overflow in *maf1*Δ leads to flux redistribution into the trehalose pathway, to protect the cells from either an increase in intracellular glucose concentration or accumulation of glycolytic intermediates downstream from glucose-6-phosphate as observed in wild-type budding yeast [77].

### Ethanol overproduction is not observed in cells lacking Maf1 during logarithmic growth

In yeast, glucose is fermented to ethanol for energy production, as it is often used as a measure of increased glycolytic flux. We examined, whether *maf1Δ* produces ethanol more efficiently than the wild type, as might be predicted from the Group 8 GO terms (Fig 2). Pyruvate decarboxylases isoenzymes Pdc5 and Pdc6 (the key enzymes in alcohol fermentation) are increased in abundance in *maf1Δ* (1.7 and 2.3 Log_2_FC respectively), suggested that Maf1 deficiency should lead to increased ethanol synthesis. We performed a fermentative capacity assay under anaerobic conditions; without cells pretreatment or with the pretreatment, when cells were glucose starved for 10 min. Under both condition there was no evidence of enhanced ethanol production in *maf1Δ* (Fig 7 C, S1 Table). Instead, accumulation of the fermentation by-product, glycerol was observed (Fig 7 D). This is consistent with increased glycolytic flux being rerouted upstream of pyruvate or downstream from acetaldehyde by the enzymes of the pyruvate dehydrogenase bypass [49].

Glycerol rather than ethanol production was also evident under aerobic conditions suggesting that access to oxygen does not affect the glycolytic flux redirection towards glycerol biosynthetic pathway in *maf1Δ*. The *maf1Δ* mutant is possibly under oxidative stress, since glycerol production has a role in response to the stress. Evidence for oxidative stress in *maf1Δ* also derives from increased protein abundance for the pentose phosphate pathway (PPP) enzymes in this mutant, which balances the systemic manifestation of reactive oxygen species and the ability to detoxify reactive intermediates [78].

### Activation of pentose phosphate pathway in cells deprived of Maf1 regulator

The comparative proteomic analysis suggests reciprocal modulation of the pentose phosphate pathway (PPP), in *rpc128-1007* and *maf1Δ* (Figs 4, 8). PPP is a source of NADPH during oxidative stress conditions. The data are consistent with an increase in flux through the PPP in *maf1Δ*, which can be achieved by increased glucose-6- phosphate dehydrogenase Zwf1 abundance, the enzyme catalyzing the rate limiting, irreversible step of the pathway. Downstream enzymes including 6-phosphogluconolactonase (Sol3, Sol4) and 6-phosphogluconate dehydrogenase Gnd2 that balance the redox potential *via* the cytosolic NADPH/NADP^+^ ratio in native yeast cells and both isoforms of transketolase (Tkl1, Tkl2) are increased in *maf1Δ*. Conversely, depletion of Zwf1, Sol4 Tkl1 and Tkl2 in *rpc128-1007* is consistent with a reduced potential of this mutant to redirect carbon flux from glucose-6-phosphate (G6P) towards 6-phosphogluconolactone (6PG) and downstream metabolic intermediates (Figs 4, 8).

**Fig 8.**
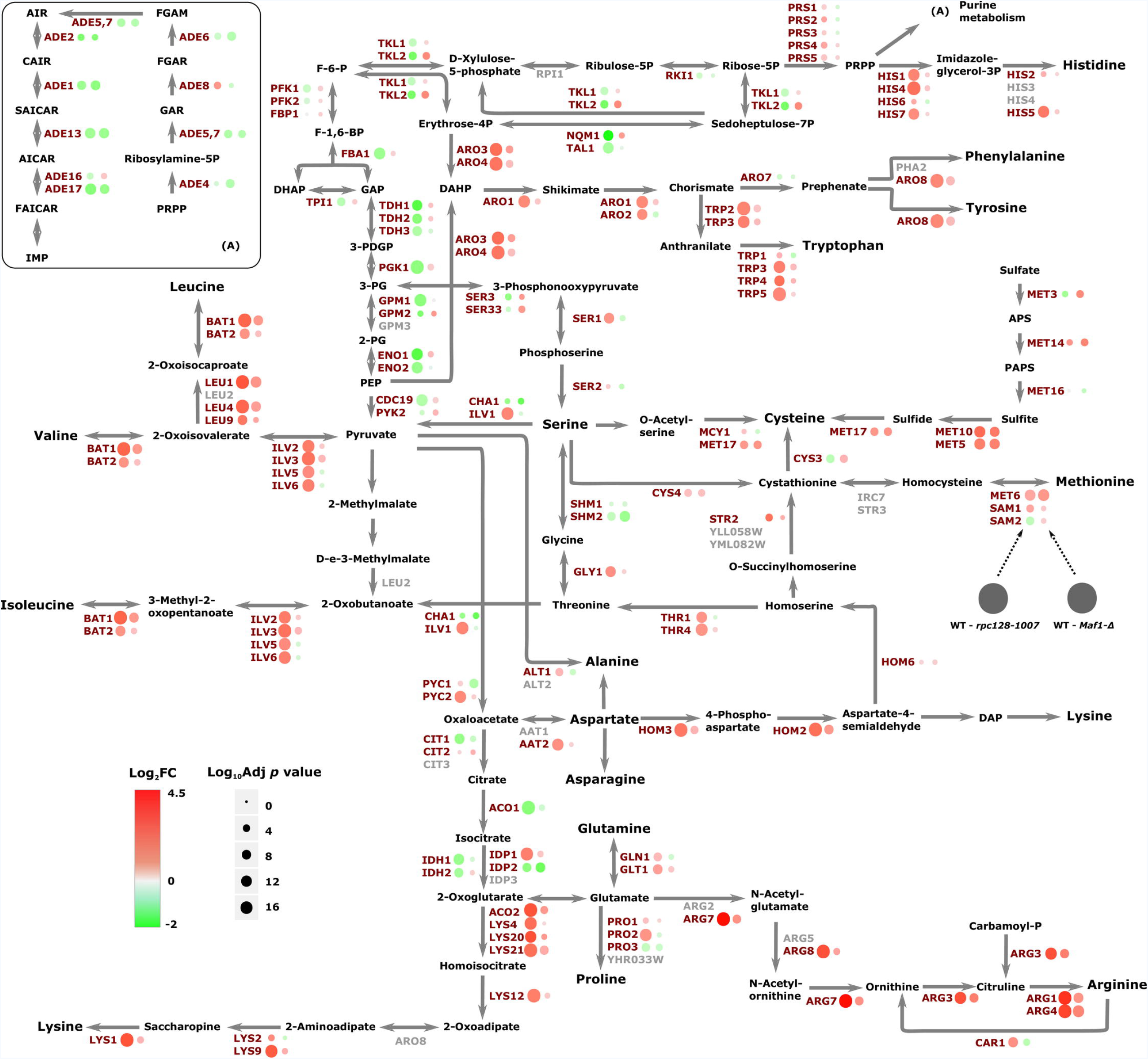
Amino acid biosynthesis and associated proteome signature. Abundance protein patterns for amino acid metabolism are presented showing those proteins with and increased abundance in red and those with an decreased abundance in green.

We grew yeast in rich medium supplemented with 2% glucose as previously and measured the Zwf1 glucose-6- phosphate dehydrogenase reaction rates (V_max_) in cell free extracts to check the potential to produce NADPH. This enzyme is highly regulated and is critical in determining the overall flow of glucose into the pentose phosphate pathway [79,80]. Zwf1 activity in *maf1Δ*, was elevated not only on glucose, as carbon source but also using glycerol whereas the activity in *rpc128-1007* remains essentially unchanged (S1 A Fig). To further corroborate the relationship between *MAF1* deletion and the oxidative stress response other proteins, such as Ctt1 stress inducible cytosolic catalase T were elevated (S2 Table). Further, total catalase activity was higher in *maf1Δ*, when compared to WT or *rpc128-1007* (S1 B Fig). The magnitude of the abundance changes in enzymes of the oxidative stress response, is observed during carbon source downshift. Here however, it occurs in the presence of glucose during the exponential phase, indicates that *MAF1* gene deletion elicits broader metabolic reprogramming than originally thought.

### RNAP III subunit *RET1*/C128 point mutation is associated with metabolic reprogramming dependent on transcriptional and translational induction by Gcn4

Glycolytic intermediates are precursors of the carbon skeletons of several amino acids. Thus, lowered glucose flux could result in the amino acid starvation response in yeast. We identified a large group of proteins that were substantially increased in abundance, that are involved in amino acid biosynthesis. The relative abundance of those was increased in *rpc128-1007* relative to WT. However, the same set of enzymes in *maf1Δ* were increased in some cases and unchanged in others. The magnitude of the increases were generally much higher in the *rpc128-1007* compared to the *maf1Δ* mutant. In *rpc128-1007*, over 30 proteins in the pathways for arginine, lysine, leucine, isoleucine, and valine biosynthesis *de novo*, along with aromatic amino acids such as histidine, tryptophan, tyrosine and threonine or their precursors were elevated (Fig 8).

In *rpc128-1007*, the decreased cellular concentration was observed in methionine biosynthesis subpathway, that is for ATP sulfurylase the product of *MET3* gene essential to catalyse the first step for assimilatory reduction of sulfate to sulfide, involved in methionine metabolism. The other proteins diminished in *rpc128- 1007* were Ser3, Ser33, Cys3 and Shm2 contributing to serine and cysteine biosynthesis.

In *maf1Δ*, the proteins Arg4, Arg3, Cpa2, Bat1, Bat2, Leu1, Leu4, Leu9 and Ilv3 were elevated; all are components of the metabolic branch that is part of L-arginine and L-leucine biosynthesis (Fig 8). By contrast *rpc128-1007*, *maf1Δ* cells exhibited enrichment in the branch of serine/cysteine, methionine biosynthesis pathway and sulfate metabolism (Met3, Met5, Met10, Met14), and these abundance changes were amongst the largest in the proteomic dataset. The expression of most genes involved in amino acid biosynthesis is under the control of the Gcn4 transcriptional activator, part of the general amino acids control (GAAC) regulon [81,82]. GCN4 transcription is stimulated by starvation for amino acids, purine, glucose limitation and specifically by initiator tRNA^met^ depletion [83,84]. Therefore, we evaluated the mRNA levels of *GCN4* by RT-PCR (Fig 9 A).

**Fig 9.**
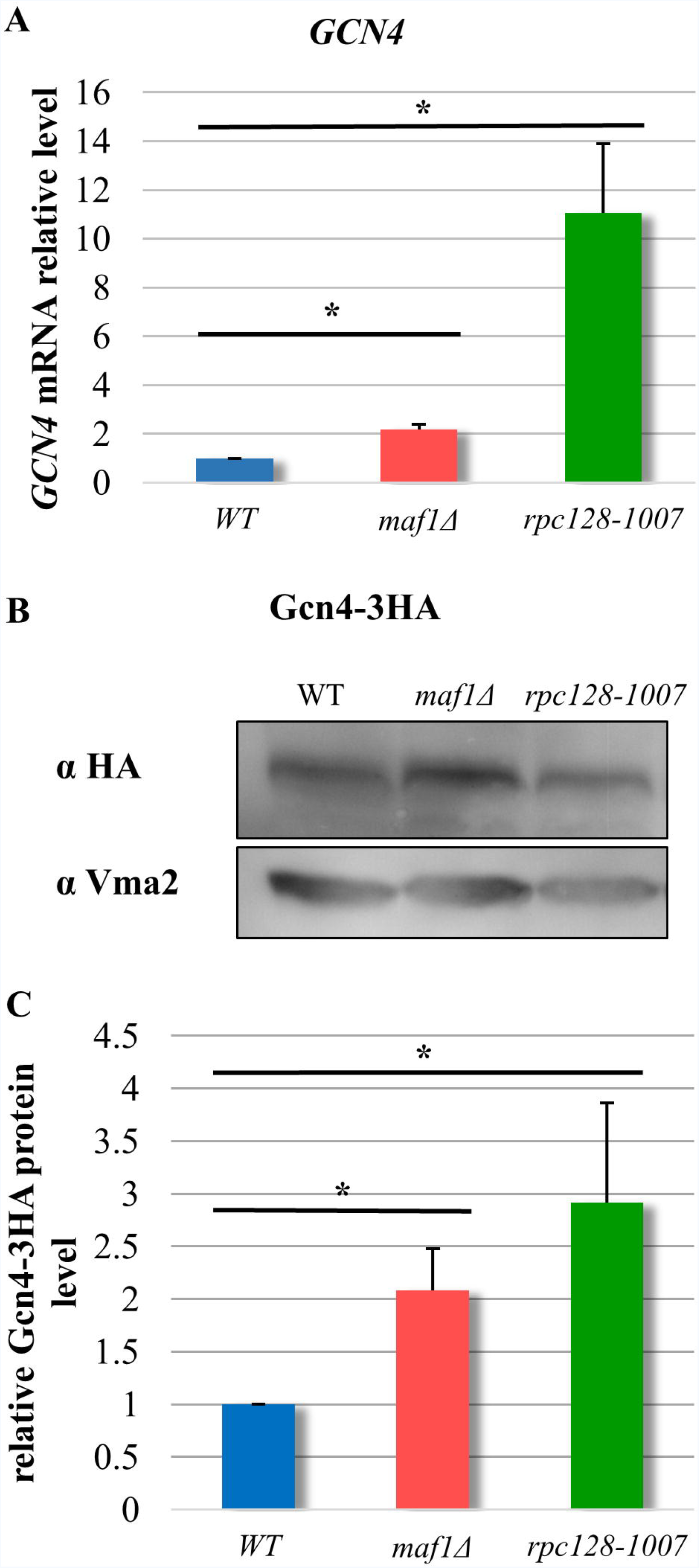
*GCN4* transcripts and Gcn4 protein relative levels are significantly increased *rpc128-1007* yeast cells and moderately in *maf1*Δ cells. Yeast cells were grown in rich medium supplemented with 2% glucose until reached exponential growth phase (A_600_ ≈ 1.0). SYBER-Green based Real-Time PCR (A) showed that *GCN4* transcript increased 2-fold in *maf1*Δ and by 11-fold in *rpc128-1007*. Wild-type strain expression level was taken as 1.0. Samples were normalized to two reference genes - *U2* spliceosomal RNA (*U2*) and small cytosolic RNA (*SCR1*). Asterisk (*) indicates *p*-values lowered than 0.05 according to t-student test. Western blotting assay (B) showed increased stability of Gcn4-3HA protein in *maf1*Δ and *rpc128-1007* mutant strains expressing chromosomally encoded Gcn4-3HA. Total cell protein extracts were subjected to SDS-PAGE and examined by Western blotting with anti-HA antibodies (B). Quantitative relative level of Gcn4–3HA protein in comparison to yeast Vma2 protein level was calculated for at least three independent biological replicates (C).

The *GCN4* mRNA steady state levels were 11-fold higher in the *rpc128-1007* and, 2-fold elevated in *maf1Δ*. The difference between these two mutants is reflected in the extent of the response in the proteomics analysis *rpc128-1007* had much stronger phenotypic change than *maf1Δ*. We constructed mutant strains with chromosomally encoded *GCN4-3HA* protein fusions to assess Gcn4 protein abundance by immunoblotting. Gcn4 abundance, normalized to Vma2 level in the total protein extracts, was elevated 3-fold in *rpc128-1007* and 2-fold in *maf1Δ*, when compared to the reference strain (Fig 9 B, C). The marked decoupling between transcript and protein changes in the strains, *rpc128-1007* (11-fold mRNA, 3-fold protein) and *maf1Δ*, (2-fold mRNA, 2-fold protein), also suggests that other regulatory factors are in operation.

For the entire proteome data set, we were able to perform transcription factor enrichment analysis, using the web-based GeneCodis tool to identify over-representation of the targets of given transcription factors in the differentially abundant proteomes (Fig 10). For *rpc128-1007*, Gcn4 was the predominant transcription factor. highlighted by the analysis for gene activation. The second most predominant transcriptional factor was Leu3 (S2 Fig) followed by Yap1 and Bas1. We also identified a group of genes with motifs enriched for GATA transcriptional factors such as Dal81, Dal80 and Gzf3 regulating genes by nitrogen catabolite repression (NCR).

**Fig 10.**
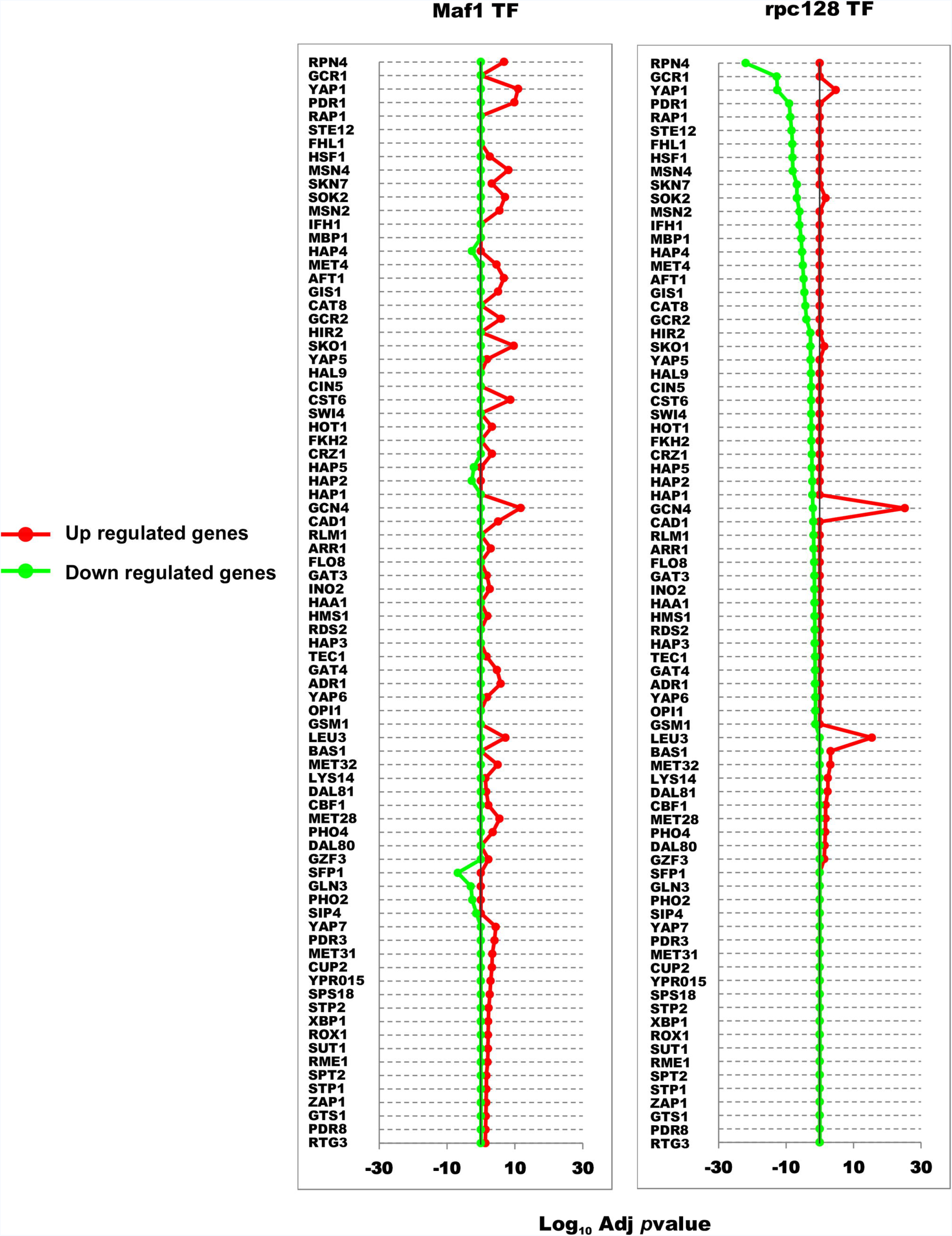
Transcription factor enrichment analysis. Enrichment in the proteome sets for individual transcription factors was calculated using the GeneCodis website taking the sets of proteins with an adjusted *p*-value of < 0.05 from both strains *maf1Δ* and *rpc128-1007* compared against the wild-type. Proteins were classified according to their positive or negative fold change and the background set consisted of all proteins identified in the given MS experiment.

The outcome of experimental and *in silico* analyses is that the dramatic increase in specific protein changes observed in *rpc128-1007* can be largely attributed to the *GCN4 stress* response and *GCN4* de-repression on glucose. By contrast to the highly focused changes in *rpc128-1007*, the gene regulatory network in *maf1Δ*, (which liberates RNAP III from regulatory circuits and nutrient signaling) (Fig 1), exhibits a broad spectrum of modest changes (including more balanced *GCN4* up-regulation) across several cellular processes, to provide the mutant with better adaptation/selective advantage to growth in glucose-rich environment.

## Discussion

Regulation of the central carbon metabolism of *S. cerevisiae* has always been a topic of considerable interest. Here, we present for the first time, the evidence that glycolytic flux in yeast can be modulated accordingly to the RNAP III activity and that central carbon metabolism adjusts to the activity of RNAP III in yeast. Our results clearly indicate the new connections between RNAP III and cellular metabolism. On the basis of the previous and the present findings, we propose that there is an internal signaling in the yeast cells, that competes against the extracellular nutrients-sensing, when a cell faces non-optimal RNAP III activity.

### An RNAP III point mutation in the *RET1*/C128 subunit is correlated with lower efficiency of glycolysis and up-regulation of *GCN4* dependent genes of amino acid metabolism

In a previous report we have shown that the *rpc128-1007* strain is insensitive to external glucose concentration cues, which manifests *via* constitutive overexpression of the *HXT2* gene, whether in glucose or glycerol growth conditions [27]. Our new data suggest that the *rpc128-1007* mutant operates under substrate limitation, such as glucose, even though the sugar is present in excess in the growth medium. RNAP III activity inhibition results in diminished abundance of enzymes in central carbon metabolism and elicits preferential synthesis of proteins involved in amino acids biosynthesis (Fig 8) attributable to the *GCN4* response. Gcn4 is translationally up-regulated in response to numerous tRNA perturbations [84,85] and yeast cells with the *RET1*/C128 point mutation produce as much as 1.6-fold less tRNA molecules compared to WT. This suppresses the defect of Maf1 inactivation, caused by increased or unbalanced levels of various tRNAs including increased tRNA^Met^ levels [13]. Initiator tRNA^Met^depletion triggers a *GCN4*-dependent reprogramming of global genome expression in response to decreased RNAP III transcription in *rpc160-112* mutant (in the largest C160 subunit) [84]. However, *HXT2* transcription is not dependent on Gcn4 transcriptional activity [84] in the *rpc160-122* mutant. *HXT2* overexpression in *rpc128-1007* is most likely related to lower glycolytic efficiency in the mutant strain [59], but the direct regulator of the phenomenon still remains to be discovered.

In the wild-type yeast cells, most glycolytic enzymes exist with significant overcapacity, regardless the carbon source. The enzymes are present in the cells at generally fixed concentration, even if reverse glycolytic processes take place [86]. However, in the *rpc128-1007* mutant, the abundance of all the glycolytic isoenzymes is reduced and concomitantly with a decrease in abundance of several proteins engaged in the side branches of the glycolytic pathway (trehalose/glycogen shunt and PP pathway). The two enzymes, fructose 1,6 bisphosphatase Fbp1 and pyruvate kinase 2 Pyk2, that catalyze the reactions in gluconeogenesis are the least affected in *rpc128-1007* in agreement with the preference of this mutant to grow on respiratory carbon sources such as glycerol [13].

In a *Drosophila melanogaster* gut model [87], reduction of RNAP III activity through controlled degradation of the C160 subunit (C160 encoded by *RPC160*) leads to diminished protein synthesis. Inhibition of RNAP III affects RNAP I, but not RNAP II-generated transcripts, suggesting that translation is the major factor regulating protein abundance in RNAP III compromised cells [87]. This further suggests that the *rpc128-1007* mutant might be limited at translation and indeed, the rpc128-1007 mutant has reduced tRNA levels [13]. In *rpc128-1007*, translation seems to be selective towards enzymes in *de novo* amino acids synthesis at the expense of the full complement of glycolytic enzymes. As a consequence of decreased abundance of the glycolytic enzymes, the glycolytic flux is very likely to decrease. Proteins could also be selectively stabilized in the *rpc128-1007* background. This accords with our finding that in *rpc128-1007* there is a decrease in RPN4 regulated elements of the protein degradation machinery, including the 26S proteasome genes [88,89] (Fig 10, S3 Table). As a result, an increase resistance to proteostatic challenge in this mutant could ensue (Fig 10). High turnover proteins may be stabilized even in an environment of diminished protein synthesis in *rpc128-1007* [90]. Overall reduction of protein catabolism might be critical for survival, consistent with a high enrichment of down-regulated proteins in this genetic background. The reduction of abundance of selected glycolytic enzymes, such as Hxk2, Tdh and Cdc19 in *rpc129-1007* is followed by the decrease in their enzymatic activity at high concentrations of glucose (Figs 5 and 6). Lowering the intracellular Cdc19 concentration is sufficient to shift from fermentative to oxidative metabolism in yeast, which reduced flux towards pyruvate [43]. The lower abundance and activity of the first (Hxk2) and the last (Cdc19) enzymes in the glycolytic pathway can be expected to reduce glycolytic flux in the mutant.

The RNAP III *RET1*/C128 mutation elicits a diminution of the low affinity glucose transporter 1 (activated on high glucose), the major glucose transporter facilitating glucose uptake under glucose rich conditions [56] and which is under control of Rgt2 (low affinity) glucose sensor (Fig 5B). External glucose signaling, may not be a dominant factor in reprogram *HXT* genes expression in *rpc128-1007*, glucose metabolism may dominate in this case [27]. Our data are consistent with the postulate that “glucose” sensing could occur intracellularly [1–4], but not yet (so far) linked to metabolic reprogramming upon change in RNAP III activity. There is some debate as to whether the signaling molecule for metabolism switching is fructose 1,6 bisphosphate [4,56]. F16BP triggers a switch in metabolism from respiration to fermentation in unicellular and higher organisms. It is the key metabolic factor determining AMPK/Snf1 kinase activity and is a potent activator of Ras pathway [3,4,64]. If F16BP plays such key roles, we reasoned that the intracellular concentration should be lower *rpc128-1007* and higher in *maf1Δ*.

Although significantly lower than in the wild-type cells, the levels of F16BP in the both mutant strains are comparable (Fig 6). This suggested that in *maf1Δ*, the mechanisms protecting cells from damaging increased concentrations of glycolytic intermediates are unperturbed thus increasing cells survival; cells capable of increased glucose consumption.

Despite the initial rise in F16BP after a glucose pulse, the metabolite attains a steady-state concentration at levels that are not likely to be toxic to the *rpc128-1007* mutant. In *rpc128-1007* cells, we presume that there is no direct relationship between intracellular F16BP and the growth defect on glucose medium exhibited by these cells, however alternative scenario is possible, if taking into account the overall abundance of the glycolytic enzymes and an excess of F16BP which may affect Ras proteins. These cells also exhibit large decrease in ribosomal proteins (RP) (S4 Table). RNAP III transcription is coordinately regulated with transcription of rDNA and ribosomal protein coding genes [91,92]. Normally ribosomal proteins and their mRNAs are stabilized when yeast is subject to increased glucose [93]. Arguably, the lower abundance of RP proteins in *rpc128-1007* could reduce the energy expense of cells that are unable to metabolize available extracellular glucose. For the *rpc128-1007*, all of the changes at the proteome level are reminiscent of the global changes in yeast cells in response to environmentally stressful, glucose deprived conditions. An exception is the cohort of proteins of the TCA cycle. RNAP III compromised cells have reduced glycolysis but this reduction does not lead to enhanced oxidative metabolism.

### *maf1Δ* cells preferentially metabolizes glucose, which results in carbon overflux fueling the side pathways dependent on glycolytic intermediates as precursors

Lack of Maf1 causes cells to reprogram their metabolism towards higher glycolytic activity when grown under high glucose conditions. This response is not reflected in an increase in abundance of the glycolytic enzymes but rather in enzymatic activity. Of course, the profile of activity modulating posttranslational modifications of these enzymes could well be different in *maf1Δ* but this was beyond the scope of this study. For example, hexokinase 2 exhibited higher activity in *maf1Δ* even though the protein abundance was reduced and the activity of this enzyme is regulated by phosphorylation [94,95]. Higher hexokinase activity should lead to increased flux into glycolysis. However, due to robustness of flux regulation, carbon is redistributed in *maf1Δ* into the side branches of the glycolytic pathway at glucose-6P (Fig 11). The *maf1Δ* shows increased capability to direct carbon into all the side branch pathways as suggested by the proteomic data and confirmed by direct metabolite assay (Fig 7 A). Glucokinase (Glk1), increased in abundance in *maf1Δ*, may redirect glucose toward glycogen storage as previously postulated [96]. The enzymes of glycogen trehalose and central carbon metabolism may be altered in *maf1Δ* as these are dependent on control by the major nutrient sensing protein kinases TOR, PKA, Snf1, Pho85 and the energy sensor Pas kinase [94,97,98].

**Fig 11.**
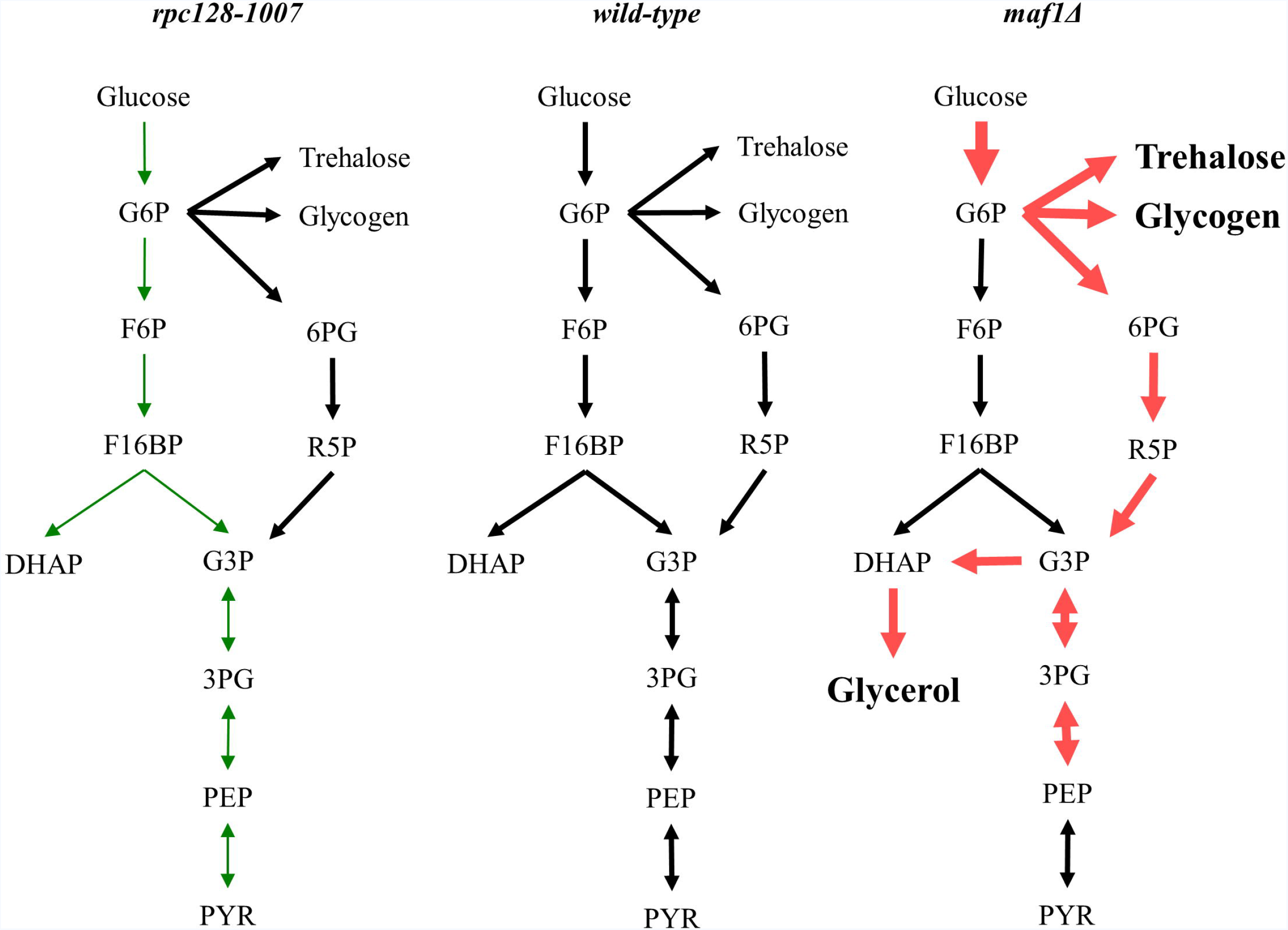
Proposed model of carbon flow in *rpc128-1007* and *maf1*Δ yeast cells. Altered RNAP III activity affects carbon flux. Low activity of RNAP III in *rpc128-1007* strain is correlated with decreased carbon flow through glycolysis in comparison to reference strain. By contrast, *maf1*Δ cells demonstrate increased carbon flow through hexokinase step and lower glycolysis compared to the control strain. In *maf1*Δ, excess glucose-6-P (G6P) is redirected into PPP and trehalose and glycogen biosynthesis. As a result fructose 1,6 bisphosphate (F16BP) concentration decreases in *maf1*Δ. From the increased glycerol concentration, carbon flux is partially redirected towards upper glycolysis at PEP. Green: decrease in carbon flux; Red: increase in carbon flux. Legend: glucose-6-phosphate (G6P); fructose-6-phosphate (F6P); fructose 1,6 bisphosphate (F16BP); dihydroxyacetone phosphate (DHAP); glyceraldehyde-3-phosphate (G3P); 6- phosphogluconate (6PG); ribose 5-phosphate (R5P); 3-phosphoglyceric acid (3PG); phosphoenolpyruvate (PEP).

In rat hepatoma cells, glucose import, and the activity of hexokinase, hexose phosphate isomerase and the glucose-6P branches that generate F16BP exert most of the flux control [99]. We believe that in *S. cerevisiae*, hexokinase 2 and increased activity of the low affinity glucose transporter Hxt1 have the greatest potential to contribute to flux rerouting in *maf1Δ*. At this stage, we do not know the intracellular signal. The activity of Hxk2 is elevated even though the trehalose shunt, which acts as safety valve against excessive supply of glucose, may correct glucose influx through allosteric inhibition of Hxk2 by trehalose-6P [72,76]. Further the glycolytic flux towards pyruvate could be enhanced by Tdh over-activation. However, carbon flux in *maf1Δ* fuel the glycerol synthesis pathway rather than causing ethanol accumulation. Glycerol accumulation in *maf1Δ* must serve as a drain for excess reducing power, to ameliorate redox imbalance in the cells*. MAF1* deletion in yeast cells is associated with redox imbalance as has previously reported by Bonhoure et al. [100] in *MAF1* knock-out mice. In *S. cerevisiae*, this is efficiently counteracted by NADH-consuming glycerol formation [101].

Activation of the branch pathways of the central carbon metabolism seen in *maf1Δ* is a hallmark of cancer cells that reprogram glycolytic activity towards synthesis of metabolites required in excess when cells rapidly divide. For instance, the PP pathway provides precursors for nucleotide and amino acid biosynthesis. This pathway, also referred to as a metabolic redox sensor, is important to maintain carbon homeostasis, is highly correlated to oncogenic, nutrient response signaling pathways [78, 102–104] and is required for NADPH regeneration. It supports metabolic reconfiguration in rapidly proliferating cells, since NADPH is a ubiquitous cofactor for most anabolic reductive reactions and got scavenging of reactive oxygen species (ROS) that cause oxidative damage to DNA and proteins and which reduce protein synthesis [105,106]. ROS scavenging enzymes also increased in abundance in *maf1Δ*, suggesting a role of Maf1 in regulation of intracellular redox potential.

How is the glucose flux distributed in Maf1 deficient yeast cells? We propose scenario, build on our observation of the decreased levels of the allosteric activator of pyruvate kinase Cdc19. The reduced F16BP concentration in *maf1Δ* cells could adversely affect Cdc19 activity. The F16BP availability for binding [67–69], but not Cdc19 abundance *per se* or phosphorylation, plays a predominant role in regulating the metabolic flux through the pyruvate kinase Cdc19 [70]. This may further lead to decrease in Cdc19 activity *in vivo* and pushing the glycolytic intermediated of lower glycolysis back to upper side branches of the pathway since lower PEP/pyruvate conversion, catalyzed by pyruvate kinase, favors accumulation of glycolytic intermediates, refueling diverging anabolic pathways, such as the pentose phosphate pathway (PPP) and serine biosynthesis [64,107].

In our model, the glycolytic flux bypasses the steps in upper glycolysis between G6P and G3P, achieved *via* improved flux through PPP. This pathway operates in three modes, depending on a cell demand for metabolic intermediates and cofactors. To avoid extensive F16BP synthesis, which would improve cells survival, the flux should be directed towards glyceraldehydes (G3P) [4]. This scenario is supported by our observation of glycerol accumulation and no change in ethanol production in *maf1*Δ cells. This glucose flux redistribution towards the PPP shunt, amino acids and nucleotide biosynthesis, due to Cdc19 action has been reported for cancer cells [108]. Further, human fibroblasts exposed to hydrogen peroxide elicit enhanced carbon flow through upper glycolysis and the oxidative branch of PPP, causing reduction in lower glycolysis activity [109].

The characteristics of *maf1Δ* in central carbon metabolism is reminiscent of cancer proliferating mammalian cells which are stimulated in the early part of glycolysis *via* PI3K/AKT activation, and making glycolytic intermediates available for macromolecular synthesis due to the low-activity isoform of PK-M2 pyruvate kinase and producing NADPH due to mutated p53 tumor suppressor [107,108].

There are further similarities between *maf1*Δ and mammalian cells after oncogenic transformation. We noted increased potential for amino acid biosynthesis, including the arginine and leucine metabolic pathways. Arg and Leu are crucial for TORC1 signaling and activation of protein translation in yeast and higher eukaryotes. Leucine is the most frequently encoded amino acid in eukaryotic genomes and its levels are sensed by leucyl-tRNA synthetase to activate TORC1 kinase [110,111]. How yeast TORC1 integrates arginine signals is presently unknown. In mammals, arginine levels are communicated by two mechanisms, involving Rag GTPases mediating amino acids signals to control mTORC1 and by a cytoplasmic mechanism that involves arginine signaling by a sensor called CASTOR [112,113].

*maf1Δ* cells have also a strong enrichment in the branch of serine/cysteine, methionine biosynthesis pathway and in sulfate metabolism by contrast with *rpc128-1007* cells. Methionine biosynthesis is connected to tRNA quality control [114]. A crucial contribution of serine/glycine to cellular metabolism is through the glycine cleavage system, which resupplies once carbon units for one-carbon metabolism The importance of serine/glycine metabolism is emphasized by genetic and functional evidence indicating that hyperactivation of the serine/glycine biosynthetic pathway drives oncogenesis. During growth on a fermentable carbon source, most serine is derived from the phosphoglycerate-3P by the gene products Ser3 and Ser33. Ser3 is a cytosolic enzyme with the dual function of phosphoglycerate dehydrogenase and alpha-ketoglutarate reductase [115]. Known for oxidizing 3-phosphoglycerate in the main serine biosynthesis pathway Ser3 also reduces alpha– ketoglutarate to D-2-hydroxyglutarate (D-2HG) using NADH, the major intracellular source of D-2HG in yeast. High levels of intracellular D-2HG are found in several types of cancer including gliomas and acute myelogenous leukemia [116].

RNAP III genes are not equally regulated by Maf1. Comparison of expression of selected tDNA genes in *maf1Δ* on glucose has revealed elevated tRNA^Met^ levels [13]. Methionine is a proteinogenic amino acid. Over-expression of initiator methionine tRNA (tRNA^Met^) leads to reprogramming of tRNA expression and increased cell metabolic activity in *Drosophila* and proliferation in human epithelial cells [117,118]. Moreover, methionine metabolism influences genomic architecture via H3K4me3 histone methylation to alter chromatin dynamics and cancer associated gene expression [119]. Furthermore, high methionine metabolism and sulfur utilisation is intertwined with high Tkl1 transketolase activity, and is dependent on the non-oxidative phase of the PPP, Tkl1 abundance increased under our study in *maf1Δ* [120]. Methionine biosynthesis, *via* the assimilation of inorganic sulfate, requires three molecules of NADPH per molecule of methionine [80]. The data presented here suggest that efficient supply of NADPH derived from the PPP in *maf1*, may also support methionine biosynthesis in the mutant.

Finally, the proteomics data obtained here with glucose grown *maf1Δ*, suggest an alternative explanation to the reduced fitness of this strain on non-fermentable carbon sources. Down-regulation of *FBP1* transcription (26) is unlikely to be the major cause of *maf1Δ* lethality when grown on glycerol. The *FBP1* gene is expressed only in the absence of glucose, and even if it is not expressed under glucose rich condition that is, under this study the Fbp1 protein is still present in *maf1Δ* at wild-type levels. It is possible that the growth defect in *maf1Δ* on glycerol is due to decrease in abundance of the enzymes involved in the glyoxylate cycle.

In conclusion, global label-free profiling of enzymatic proteins in yeast has provided new insight in metabolic physiology. Protein abundance patterns characterized in the mutant strains that show different phenotypes on fermentable and non-fermentable carbon source highlighted metabolic pathways that could now be the target for further genetic or metabolic analysis. This work emphasises *S. cerevisiae* as a very good model organism for systems level studies on the dynamics of cellular networks. There is growing evidence for contradictory observations in cultured human cancer cells and in multicellular organisms including mouse models [100,119]. We anticipate that yeast cells will continue to be appreciated as a source of basic biological information in building an integrated picture of metabolism and gene regulation.

## Conclusions

- The capacity of the glycolytic pathway can be altered by manipulation of RNAP III activity.
- Lack of Maf1, the negative regulator of RNAP III driven non-coding RNA transcription, enhances glycolytic flux and results in accumulation of end products upstream of pyruvate.
- Severe reduction in growth rate caused by RNAP III mutation *rpc128-1007* on glucose is correlated with a decrease in abundance of glycolytic enzymes.
- The translation machinery in the *rpc128-1007* mutant seems to be selective towards mRNA coding for enzymes for amino acids synthesis *de novo* at the expense of full complement of glycolytic enzymes.
- The critical decrease in abundance of glyoxylate cycle enzymes reduced the ability to convert non-fermentable substrates in *MAF1* knockdown yeast.

## Acknowledgments

We want to thank Mark Johnston (Dept. of Biochemistry and Molecular Genetics, University of Colorado Denver, US) for providing *pBM2636* plasmid and Emil Furmanek for preparation of samples for proteomics analysis. We are grateful to Dr Philip Brownridge for excellent instrument support. This work was supported by National Science Centre, Poland grant 2012/05/E/NZ2/00583 to M.A. and by funding from Faculty of Chemistry, Warsaw University of Technology, Poland. I dedicate this work to my baby son Piotr.

## Supporting information

**S1 Fig. Zwf1 (A) and the Ctt1 catalase activity (B) is increased in Maf1 deficient mutant.**

Yeast cells logarithmically growing in YPD 2% glucose medium were harvested at A_600_ ≈ 1.0. For Zwf1 activity assay, NADH breakdown was measured at 340 nm in time at 30°C. For catalase activity, hydrogen peroxide decomposition in reaction mixtures containing yeast cell free extracts was monitored as change in absorbance at 240 nm in time at 30°C. Results are presented as total mean enzymatic activity from five independent biological replicates with standard deviation expressed as μmol·min^-1^·mg^-1^ protein. Asterix (*) indicate *p*-value < 0.05 according to t-student test.

**S2 Fig. Real-time quantitative PCR analyses of *LEU3* gene transcript.**

Slight increase in *LEU3* transcript levels was observed in *rpc128-1007* in glucose medium. Opposite effect was observed on glycerol-based rich medium. Maf1 deficiency cause 2-fold decrease in *LEU3* mRNA levels, when grown non-fermentative carbon source and reduced in glucose-based rich medium. Yeast cells were grown in 2% glucose (YPD) or 2% glycerol (YPGly) rich medium at 30°C until an A_600_ ≈ 1.0. SYBER-Green based Real-Time PCR was performed. The expression level of each target PCR product was normalized to reference genes transcript levels: *U2* spliceosomal RNA (*U2*) and small cytosolic RNA (*SCR1*). The means + standard deviations of the relative expression levels from three independent biological replicates are shown. The value of basal gene expression level of WT strain was assumed as 1.0. Asterisks (*) indicate p-values ≤ 0.05 determined by t-student test.

**S1 Table. Mean value for triplicate experiments of fermentative capacity assay of WT and *maf1*Δ cells.**

Results are expressed in mM / g dry weight. ‘C-limited’ stands for ‘carbon-limited conditions’.

**S2 Table. The list of 8 proteins involved in oxidative stress with higher abundance in *maf1*Δ and lowered quantity in *rpc128-1007* mutant in comparison to wild-type reference strain under high glucose conditions according to global proteomics.**

**S3 Table. The list of 18 proteins regulated by RPN4 that are part of 26S proteasome and its core complex, 20S proteasome, with lowered abundance in *rpc128-1007* mutant in comparison to wild-type reference strain under high glucose conditions according to global proteomics.**

**S4 Table. The list of 73 proteins involved in ribosome biogenesis with lowered abundance in *rpc128-1007* mutant in comparison to wild-type reference strain under high glucose conditions according to global proteomics.**

